# Differentiable Design for Morphogenesis I: Simulation and Simulacra

**DOI:** 10.64898/2026.07.02.736195

**Authors:** Ozgur Beker, Bianca Dumitrascu

## Abstract

Cells build tissues through local exchanges of force and information, yet the rules governing these interactions are difficult to infer from sparse observations. Here, we introduce *waxMorph*, a differentiable cell-based framework for generating and reconstructing three-dimensional morphogenesis. In synthetic and biological data, *waxMorph* reproduced established mechanochemical shape programs, inferred continuous trajectories from static tissue volumes, and recovered spatially organized latent signals. In a developing mouse myocardium dataset, it reconstructed unobserved intermediate geometries more accurately than optimal-transport interpolation, while in forelimbs it distinguished related developmental trajectories. By varying the capacity and spatial organization of the latent cues available to cells, *waxMorph* also provides a model-based way to quantify the complexity of shape assembly. *waxMorph* is built within the spatial-computing ecosystem of NVIDIA Warp. It provides an open-source, Python-native, GPU-accelerated, hybrid physics–AI framework for learning how local cellular interactions give rise to biological form.

## Introduction

Morphogenesis is the making of biological form [1]. In development, form emerges iteratively from the coordinated actions of cells. As their molecular states change, cells signal to one another and generate forces that reshape the tissue. Through these actions, tissues fold, elongate, branch, and eventually assemble into functional organs. Understanding morphogenesis requires multiscale models that connect molecular and mechanical rules at the cell level to tissue-scale deformations.

Questions about biological form date back at least to Aristotle [2–4]. Computational models of morphogenesis are comparatively recent. This tradition began with attempts to relate biological shape to growth, geometry, and physical constraints [1], and later expanded to formal theories of chemical patterning [5, 6]. Turing showed that local reaction and diffusion could generate spatial order from initially homogeneous states [5]. His model provided one of the first mathematical accounts of self-organized biological patterning. Later, positional information frameworks proposed that cells interpret morphogen concentrations as molecular coordinates that guide fate decisions [7–10]. These ideas made pattern formation a central topic in developmental biology [11, 12].

Form, however, is more than a pattern of massless points. To make a fold, tube, branch, or cavity, cells must carry and sculpt material through space and time [13]. Recent computational approaches account for the added complexity by treating morphogenesis as a mechanical process. In this view, tissues are active materials remodeled by cellular forces, growth and movement. Continuum models describe tissues as deforming sheets, fluids, or elastic bodies, often using partial differential equations to model velocity, stress, density, or molecular fields on changing domains [14–21]. Continuum perspectives have led to concepts such as morphogenetic flow and dynamic morphoskeletons [14]. These use ideas from fluid mechanics and Lagrangian coherent structures to describe tissue deformation through time [14, 22]. By contrast, vertex, cellular Potts, spheroid, and agent-based models represent tissues as collections of interacting cells [23–29]. These approaches have shown how local rules can generate global deformation, and how molecular patterning and mechanical forces can be coupled in the same developmental process. They have also highlighted that morphogenesis cannot be assigned to one level of description, as it is a joint consequence of molecular state, cellular behavior, tissue geometry, and physical constraints.

Most computational models of morphogenesis are forward models. Researchers specify initial conditions and local interaction rules, then observe the spatiotemporal dynamics and morphologies that emerge. These trajectories aim to capture key features of real data. In vertex models for example, cells are represented as polygons whose vertices move to minimize a global energy function. Typically, the energy function contains terms encoding mechanical properties such as edge contractility or area and perimeter elasticity [30–32]. By varying these parameters, these models reproduce tissue-scale behaviors such as collective flows, cell rearrangements, and large-scale deformations [33, 34].

Advances in differentiable programming and generative modeling have made inverse problems in tissue mechanics increasingly tractable [35]. New differentiable vertex models cast epithelial mechanics, parameter inference, and inverse mechanical design as bilevel optimization problems [36]. This is a powerful formulation for learning mechanics within a specified model, especially for two-dimensional confluent epithelia. However, the mechanical constraints are largely fixed in advance, molecular state is not explicitly coupled to tissue deformation, and the central object of inference is mechanical parameterization rather than the local rules of shape assembly.

A complementary line of work uses differentiable programming to design collective cellular systems. Neural cellular automata can grow, maintain, and regenerate target patterns [37, 38]. Differentiable cell-based simulations can optimize genetic and interaction parameters for symmetry breaking, growth control, and pattern formation in two dimensions [39]. Related graph-based methods use equivariant neural networks to move point-like agents toward target three-dimensional shapes [40]. Together, these studies show that learned interactions can generate coordinated multicellular dynamics and assemble prescribed forms. They do not, however, address the inverse problem we study: inferring three-dimensional developmental trajectories from observed tissue volumes while representing volumetric cells, changing neighborhoods, and explicit physical and mechanochemical constraints.

Here we introduce *waxMorph*, a differentiable framework that unifies simulation and inference for three-dimensional morphogenesis using spheroidal cell agents. Given specified mechanochemical rules, *waxMorph* simulates the tissue trajectories they produce. Given observed morphologies, it learns neighbor-dependent cell-state updates that connect them while enforcing spatial adjacency, adhesion, volume exclusion, and molecular diffusion. Implemented in Python with NVIDIA Warp, PyTorch, and JAX, *waxMorph* supports differentiable simulation, learned deformation, visualization, and downstream analysis within a shared cellular representation [41–43]. We evaluate the framework on established mechanochemical growth programs, synthetic shape transformations, developing mouse forelimbs, and mouse myocardium trajectories [27, 44, 45].

## Results

### A hybrid physics–deep learning ecosystem for in silico morphogenesis

Cells are fundamental units of computation. To understand or build systems that recapitulate morphogenesis, we first need to specify what an individual cell can sense and do. At minimum, cells need bodies that occupy space and move through it, internal states that encode their intrinsic molecular properties, sensors that can read in information from their environments, and rules that determine how they respond to these cues. In turn, these responses can alter both cells and their neighborhoods by coupling local cellular decisions to global changes reflected in tissue shapes. Our in silico morphogenesis ecosystem, *waxMorph*, is developed around this minimal representation. We model each cell as a three-dimensional agent with position, volume, polarity, molecular state, and a dynamically changing set of neighbors (Fig. 1, left). Cells sense local molecular and mechanical cues and respond by changing state, moving, growing, or dividing.

**Figure 1:**
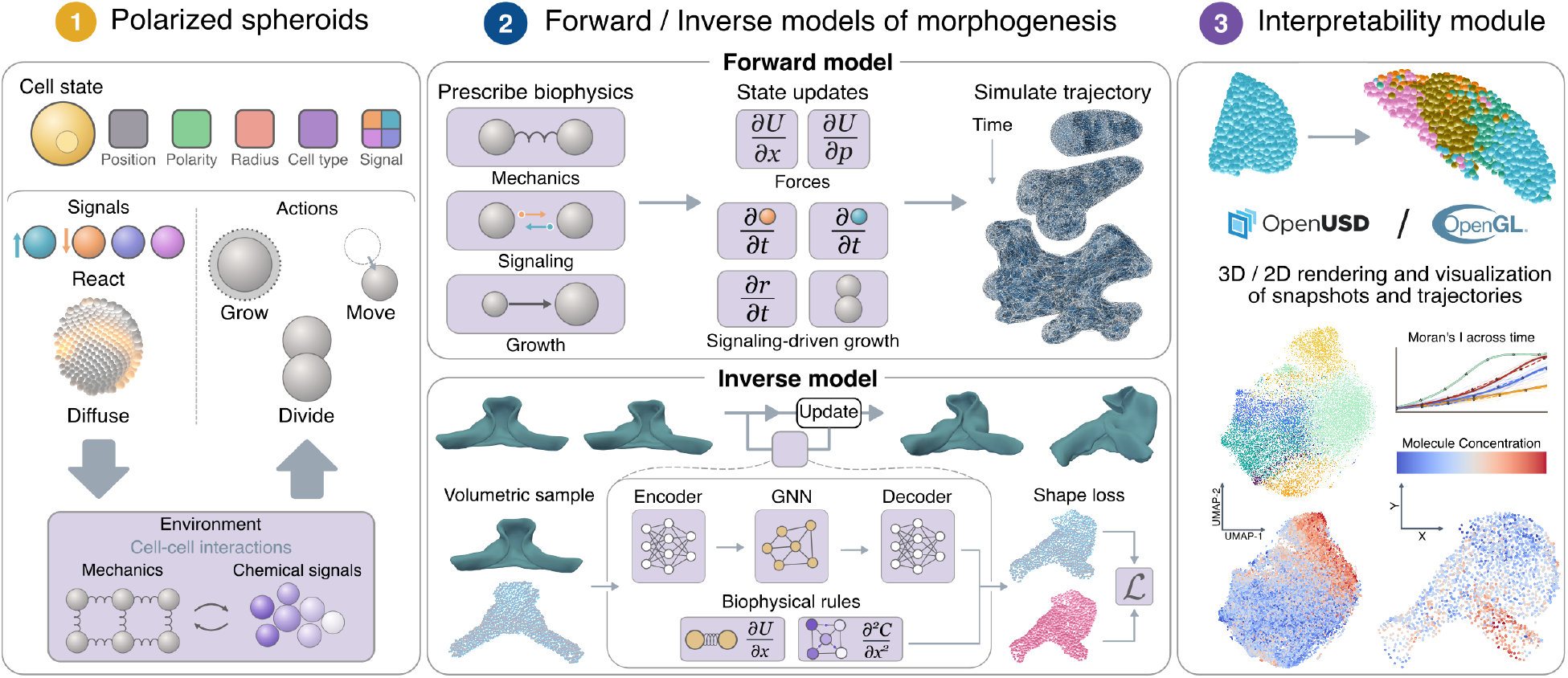
*waxMorph* integrates differentiable mechanochemical simulation, learned tissue deformation, and trajectory analysis. **1: Polarized spheroids**. Cells are represented as three-dimensional spheroids with position, volume, polarity, molecular state. Local molecular and mechanical interactions update individual cell states and collectively reshape the tissue. **2, top: A Forward model of morphogenesis**. Users prescribe molecular dynamics and mechanical potentials. Automatic differentiation converts the specified potentials into force and state updates, enabling differentiable simulation of growth, division, signaling, and tissue mechanics. **2, bottom: An inverse model of multicellular deformations**. A graph neural network simulator learns neighbor-dependent updates to cell position, polarity, and latent signaling state from observed tissue shapes. Message passing is restricted to spatially adjacent cells, while prescribed physical constraints enforce volume exclusion, short-range adhesion, and diffusion. **3: Interpretability Module**. Simulated and learned trajectories can be rendered as two- or three-dimensional images and videos or exported as Universal Scene Description files for downstream visualization. The resulting cellular trajectories can be analyzed using conventional single-cell and spatial-analysis workflows or as densely sampled functions of developmental time.

This representation supports two complementary modes. For forward simulation, users specify molecular dynamics and mechanical potentials governing interactions between neighboring cells. Kernel-based automatic differentiation transforms these potentials into cellular forces, allowing mechanical interactions to be modified or combined without deriving each force law analytically (Fig. 1, middle, top) [41].

For inverse design, a graph neural network simulator learns individual cell updates from information shared with its spatial neighbors. Node and edge features of a changing spatial graph are used to predict updates to cell position, polarity, and molecular state. Message passing is restricted to adjacent cells, soft-sphere interactions enforce volume exclusion and short-range adhesion, and signaling molecules diffuse between neighbors (Fig. 1, middle, bottom). A permutation-invariant shape loss and trajectory-smoothing penalty allow the model to infer deformations from three-dimensional tissue meshes, including volumes reconstructed by Optical Projection Tomography (OPT) or from two-dimensional segmentations [44, 45].

Finally, we render the learned trajectories as three-dimensional cellular tissues and export their molecular, mechanical, and geometric states for downstream analysis (Fig. 1, right). Complete scene descriptions are stored using the Universal Scene Description (USD) format. USD is an open source file type originally developed by Pixar Animation Studios, compatible with modern visualization tools [46, 47]. The same outputs can be analyzed using spatial statistics, single-cell workflows, and functional data analysis. Together, these components form an end-to-end hybrid physics–deep learning ecosystem in which a shared cellular representation supports forward simulation, inverse design, visualization, and analysis.

### A minimal mechanochemical circuit generates distinct three-dimensional shape programs

*waxMorph* represents each cell *i* as a spherical agent with a cell state **s**_*i*_, encoding its position **x**_*i*_, polarity **p**_*i*_, radius *r*_*i*_, equilibrium radius 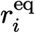, and molecular concentrations **c**_*i*_. We use this representation to ask whether Turing spheroids can reproduce the patterning and growth behaviours of the three-dimensional Turing vertex model [25–27, 48].

We prescribe a two-component activator–inhibitor circuit in which the activator promotes its own production and induces an inhibitor that restrains activator accumulation (Fig. 2A; *Methods*) [5, 27, 49]. The activator diffuses more slowly than the inhibitor, allowing initially weak molecular differences to develop into periodically spaced activated regions. The spatial characteristic *χ* specifies the activator diffusivity relative to that of the inhibitor, whereas the reaction rate *γ* sets the timescale of the reaction–diffusion dynamics. We embed this circuit in an epithelial–mesenchymal aggregate comprising a polarized epithelial shell surrounding a softer mesenchymal interior (Fig. 2B) [28, 50]. Mechanical potentials orient epithelial polarity perpendicular to the local sheet and suppress stacking of epithelial cells, maintaining an enclosed surface around the growing mesenchyme. Cell positions determine which neighbors exchange activator and inhibitor molecules. In turn, the intracellular activator concentration regulates mesenchymal growth and division, altering cell positions and rebuilding the signaling graph. The tissue therefore couples molecular patterning and mechanics through its changing spatial organization.

**Figure 2:**
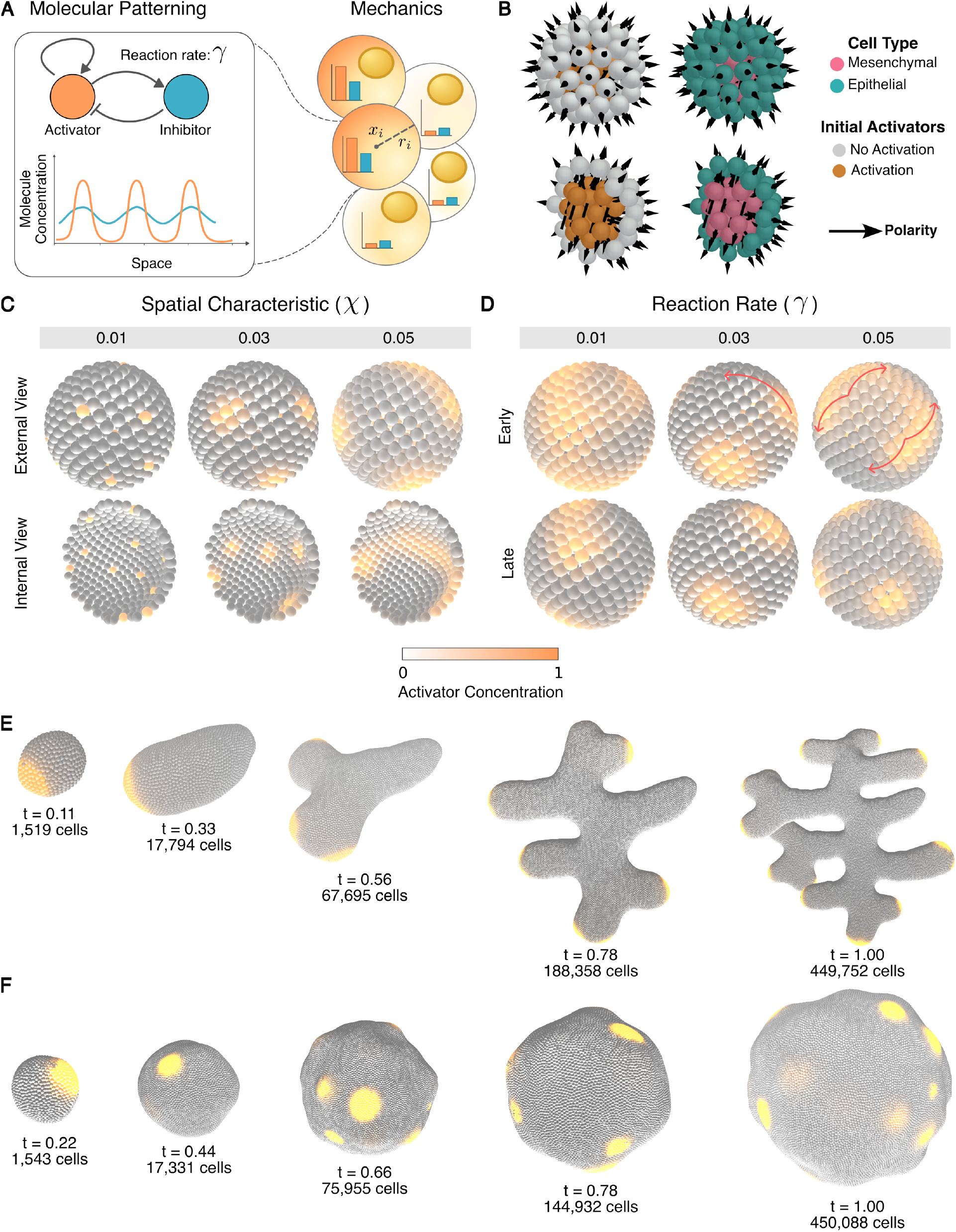
A minimal activator–inhibitor circuit generates distinct three-dimensional morphogenetic programs. **A:** Two-component reaction–diffusion circuit case study. The slowly diffusing activator promotes its own production and induces a more rapidly diffusing inhibitor with periodically spaced activation domains. **B:** Surface and cross-sectional views of the initialized epithelial–mesenchymal aggregate. A polarized epithelial monolayer encloses a softer mesenchymal population in which activator concentration controls growth and division. Colors denote cell type and activator state. **C:** Increasing the spatial characteristic *χ* enlarges the activated domains; sufficiently large values destabilize spatial patterning. **D:** A low reaction rate *γ* produces stable activation domains, whereas higher rates generate domains that form and move more rapidly. **E:** Low reaction rates maintain localized growth domains that break spherical symmetry and produce elongation and branching. **F:** High reaction rates redistribute activation across the tissue surface producing repeated surface undulations.

We first examine molecular patterning in a tissue of fixed size. Increasing *χ* enlarges the activated regions formed across the tissue (Fig. 2C). At sufficiently large values, the relative diffusivities of the activator and inhibitor no longer support a stable spatial pattern. The system instead undergoes repeated activator–inhibitor pulses before converging to a state in which patterning is lost (Supp. Fig. S2) [5, 49]. Changing *γ* alters how rapidly the pattern develops. Low reaction rates produce activated regions that remain comparatively stable over the simulated interval, whereas higher reaction rates produce regions that establish and move more rapidly (Fig. 2D).

When cells are allowed to grow, *waxMorph* recapitulates the elongation–branching and surface-undulation regimes reported in the three-dimensional Turing vertex model [27]. At low reaction rates, stable activator domains increase the equilibrium radius of nearby mesenchymal cells, driving local growth and division. The aggregate then breaks spherical symmetry and elongates along the activated regions (Fig. 2E). As the tissue grows, activated regions divide into separate domains, establishing additional growth axes and producing branching. At high reaction rates, activated regions move across the tissue before a single region can sustain a dominant growth axis. Growth is redistributed over the surface, producing a rounded aggregate with repeated surface undulations (Fig. 2F). These results show that *waxMorph* recovers the molecular and morphological regimes of the three-dimensional Turing vertex model without relying on its rigid vertex-based topology. Instead, the same shapes emerge from volumetric spheroidal cells with dynamically changing neighbors, allowing tissues to be represented as filled aggregates.

### Automatic differentiation preserves morphogenetic shape grammar

To generate forward simulations, *waxMorph* uses automatic differentiation to obtain forces from mechanical potentials without deriving each force law analytically. This makes the mechanical model modular: physical interactions can be added, removed, or modified by changing the potential, while the corresponding force updates are generated automatically. Automatic differentiation also reduces algebraic and implementation errors and preserves a differentiable path through the simulator for downstream optimization.

To test the quality of morphologies produced by automatic differentiation, we compare them to morphologies derived from analytical force updates in two regimes: elongation and branching at low reaction rates, and surface undulation at high reaction rates (Fig. 3A; *Methods*). For each regime, we generated trajectories using either analytically derived gradients or automatic differentiation of the same mechanical potentials (Fig. 3B; Appendix A). We simulated 20 replicates per condition, grew each tissue to 80, 000 cells, and aligned the resulting three-dimensional shapes for translation and rotation before comparison (*Methods*).

**Figure 3:**
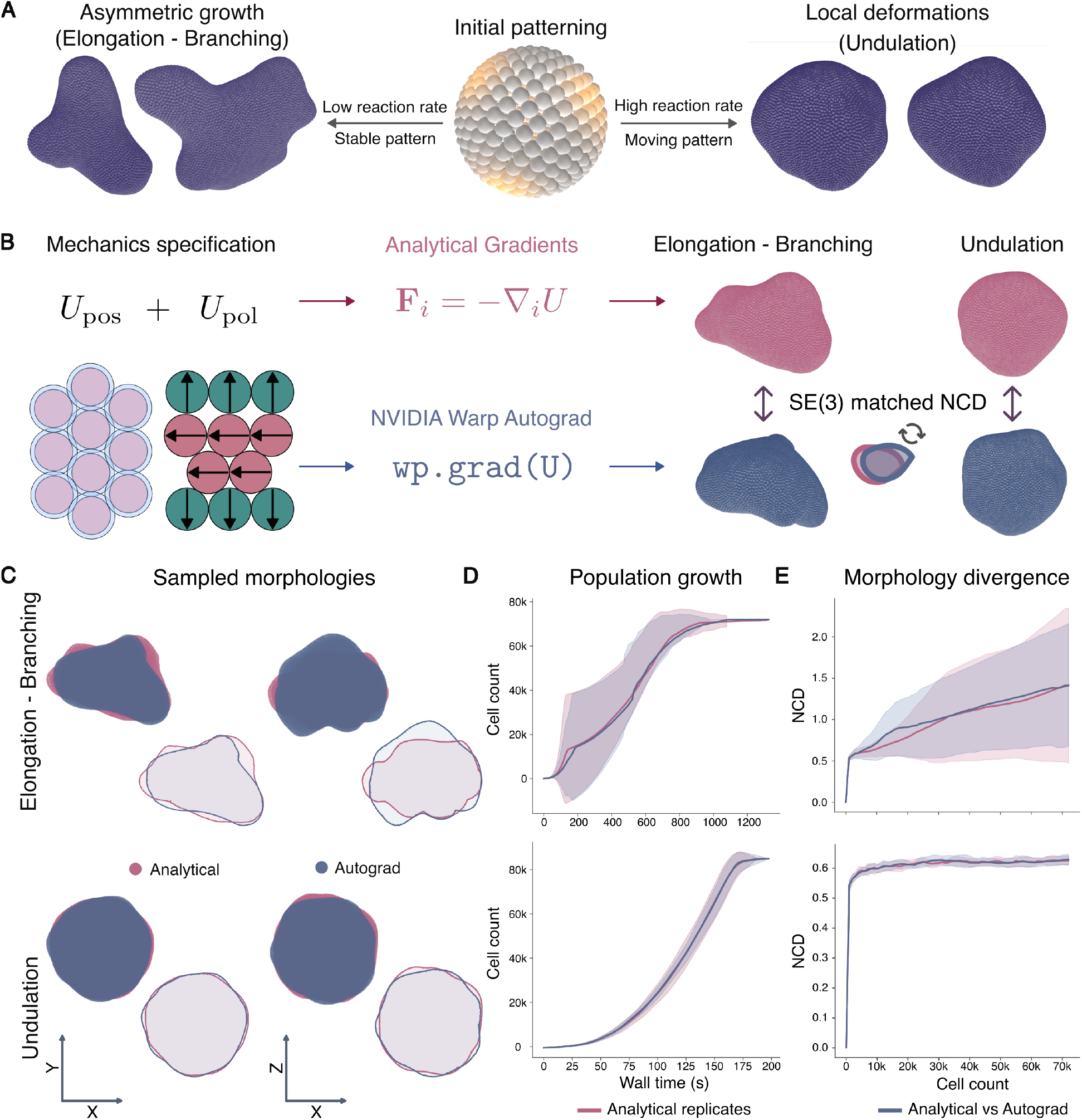
Automatic differentiation preserves the morphogenetic programs encoded by mechanical potentials. **A:** Two mechanochemical regimes used to compare analytically derived force updates with forces obtained by automatic differentiation: elongation and branching at low reaction rates and surface undulation at high reaction rates. **B:** Experimental design. Updates derived from closed-form analytical gradients and automatic differentiation of the mechanical potential were compared across twenty independent replicates. Per replicate, tissues were simulated until they reached 80,000 cells. Distance between shapes was computed using the rotation / translation matched normalized Chamfer distance (NCD), which is the average distance between nearest neighbors across volumes. **C:** Representative final morphologies from analytical and automatically differentiated simulations after alignment for translation and rotation. Examples show the closest-matching replicate pair for each regime. **D:** Tissue growth trajectories produced by analytical and automatically differentiated force updates. Comparisons are shown between independent analytical replicates and between analytical and automatically differentiated replicates. **E:** Morphological differences were measured at increments of 1,000 cells using the normalized Chamfer distance post rigid alignment. Analytical–automatic-differentiation differences remain comparable to the variation between independent analytical simulations.

Automatic differentiation recovered the characteristic morphologies of both regimes (Fig. 3C). The two implementations followed similar growth trajectories, including in the elongation–branching regime, where replicate-to-replicate variation was greater (Fig. 3D). We quantified this agreement by measuring shape divergence after every increase of 1, 000 cells. Across the full simulation trajectory, differences between analytical and automatically differentiated simulations remained comparable to those between independent analytical replicates (Fig. 3E). Thus, automatic differentiation can replace hand-derived force updates without changing the target shape trajectory, making the simulator easier to extend and optimize.

### *waxMorph* learns biophysically constrained tissue deformations

The inverse-learning module of *waxMorph* learns cell-state updates leading to observed 3D tissue shapes. Given an initial state, a target state, a signaling network, and a set of physical constraints, *waxMorph* generates a plausible trajectory between the two snapshots (Fig. 4A, left). When intermediate snapshots are available, the learned trajectory passes through all observed states (Fig. 4A, right).

**Figure 4:**
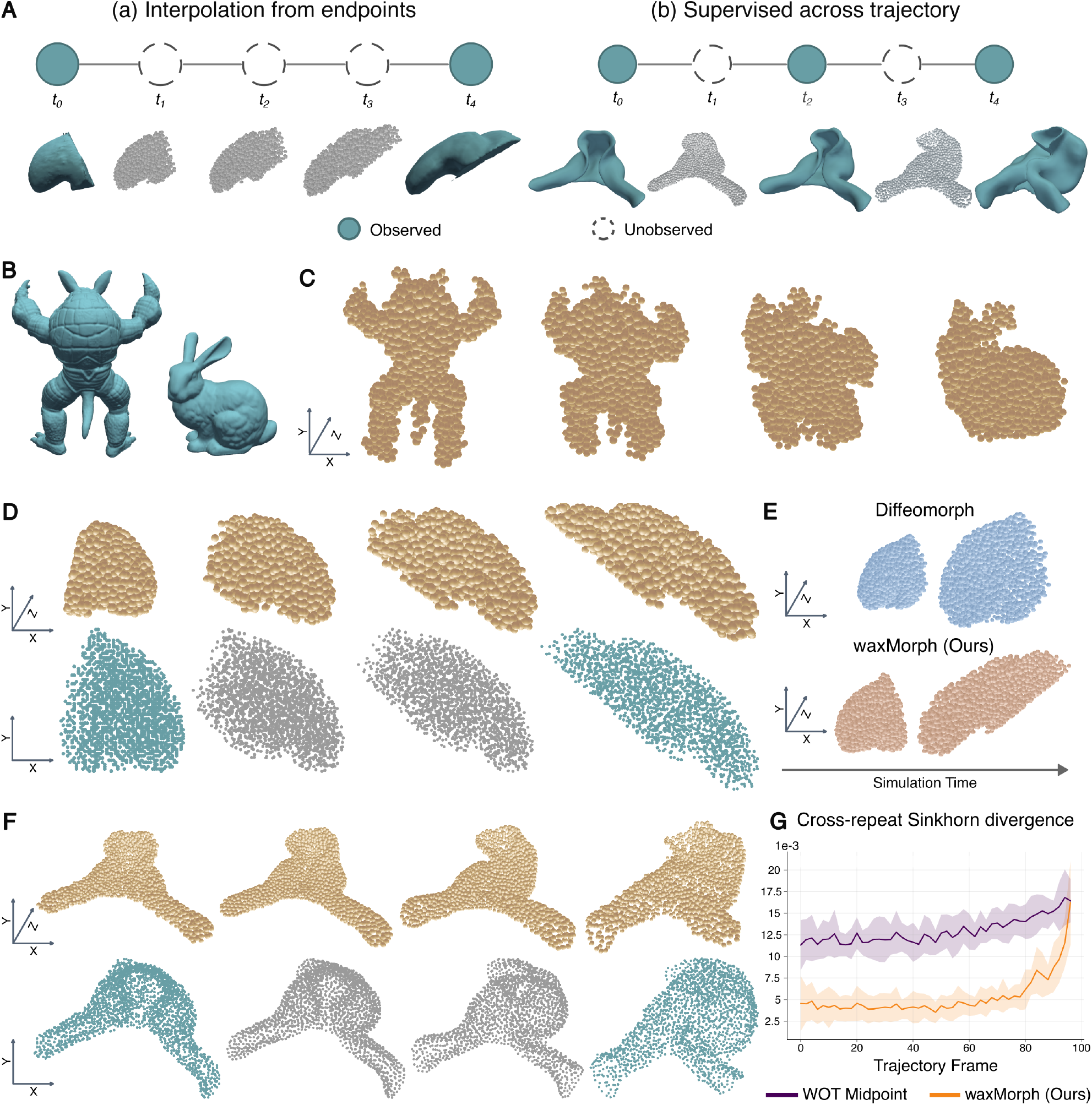
Biophysical constraints support reconstruction of synthetic and developmental shape trajectories. **A:** Inverse-learning formulation. Given an initial tissue state and one or more observed target shapes, a graph neural network learns local cell-state updates that connect the observations while enforcing physical and molecular constraints. **B:** Armadillo and Stanford Bunny meshes used as a synthetic large-deformation benchmark. Mesh interiors were sampled with 2,000 spheroidal agents. **C**, Learned Armadillo-to-Bunny trajectory. Local neighbor-dependent updates transform the initial cell population into the target while preserving non-overlapping cell volumes. **D:** Learned early-to-mid mouse forelimb trajectory from Optical Projection Tomography data, shown as three-dimensional renderings (top) and projections of cell centers onto the *xy* plane (bottom) [44]. **E:** Endpoint reconstruction by *waxMorph* and DiffeoMorph for the same forelimb transformation. **F:** Reconstruction of a densely sampled mouse myocardium trajectory, shown as three-dimensional renderings (top) and *xy* projections of cell centers (bottom) [45]. Every other observed frame was withheld during training. **G:** Reconstruction error for the held-out myocardium frames using *waxMorph* or midpoint interpolation based on Waddington optimal transport (WOT). *waxMorph* yields lower error across the reconstructed trajectory.

Cells interact through a dynamic spatial graph, with message passing restricted to adjacent neighbors. Volume exclusion and short-range adhesion are enforced through a soft-sphere potential, and latent signaling molecules diffuse across the neighborhood graph (Supp. Fig. S3A,B). Reciprocal mechanical interactions are applied explicitly, whereas a graph neural network learns the remaining neighbor-dependent updates to cell position, polarity, and latent signaling state [51]. Diffusion promotes locally correlated signaling patterns, which coordinate assembly of the target morphology. The model is trained with a permutation-invariant shape-matching loss between predicted and target cell distributions. We encourage smooth deformations with a regularization term penalizing large position changes between consecutive frames.

We asked if *waxMorph* could learn the local cell-state updates needed to generate a large deformation between shapes with distinct geometry (Fig. 4B). We applied *waxMorph* to two synthetic meshes widely used in computer graphics: the ‘Armadillo’ and the ‘Stanford Bunny’ [52, 53]. Since the meshes don’t carry single-cell morphologies, we used Poisson-disk sampling to fill each mesh with a dense, evenly spaced, and connected population of spherical agents (Supp. Fig. S4; *Methods*) [54–56]. *waxMorph* successfully learned local updates that transformed the Armadillo into the Bunny while satisfying the prescribed physical constraints and preserving spatial neighborhood contiguity (Fig. 4C).

We next interpolated between the early and mid stages of a previously published mouse forelimb development dataset (Fig. 4D) [44]. We benchmark our method against DiffeoMorph, a physics-free, point-cloud based method [40]. DiffeoMorph failed to converge to the correct target morphology. In contrast, *waxMorph* more closely recovered the target limb shape throughout the trajectory. Due to its explicit physical constrains, *waxMorph* also maintained the nonoverlapping volumes of individual cells (Fig. 4E, Supp. Fig. S5A).

We further tested whether *waxMorph* could recover intermediate tissue shapes from sparsely observed developmental states. We used a published pseudodynamic atlas of mouse heart-tube morphogenesis [45], from which we sampled 100 frames spanning the later stages of heart-tube morphogenesis. We held out every other frame during training (Fig. 4F, Supp. Fig. S5B). We compared the reconstructed frames against midpoint interpolations obtained with WaddingtonOT [57]. As WaddingtonOT is an optimal transport technique originally designed for the analysis of single-cell data, we first modified it to work with point-cloud inputs. *waxMorph* achieved significantly lower error throughout the trajectory, indicating its flexibility in modeling physical interactions throughout densely sampled trajectories (Fig. 4G).

### Latent molecular states organize deformation across space and time

In the forward module, molecular concentrations are specified as part of the model. Activator and inhibitor dynamics determine which spatial patterns form and where growth occurs. In biological data, however, three-dimensional tissue geometries are rarely accompanied by matched molecular measurements. Live imaging can capture morphology over time, but only a limited number of genes can typically be labeled, and full spatial transcriptomes are usually unavailable. We therefore ask whether the signals needed to coordinate cellular movement can be inferred directly from data. In the inverse module, these signals are represented as latent molecular states that cells use to communicate and assemble the target morphology. Each cell is assigned a latent molecular signaling state, **c**_*i*_, which is shared with its spatial neighbors and learned during emulation.

We first learn the signals required to achieve the deformation of the Armadillo volume into the Stanford Bunny (Fig. 5A). As the volumes changed shape, cells with similar latent states formed spatial domains. At the final frame, these domains remained visible in both the physical space and the two-dimensional UMAP [58] space of the latent molecular states (Fig. 5B). At the final Bunny morphology, cells with similar latent states formed coherent spatial domains corresponding broadly to anatomical regions such as the head, torso, and legs (Fig. 5B). We quantified positional information as the mutual information between latent states and cell positions, following the positional information framework [7, 8]. Mutual information was estimated for both the continuous latent-state vectors and their Leiden cluster assignments using the KSG estimator [59]. We separately measured the spatial autocorrelation of each latent signal using global Moran’s *I* [60]. The latent signals showed significant spatial autocorrelation under permutation testing (*p <* 0.001) and were therefore both positionally informative and spatially organized (Fig. 5C; *Methods*).

**Figure 5:**
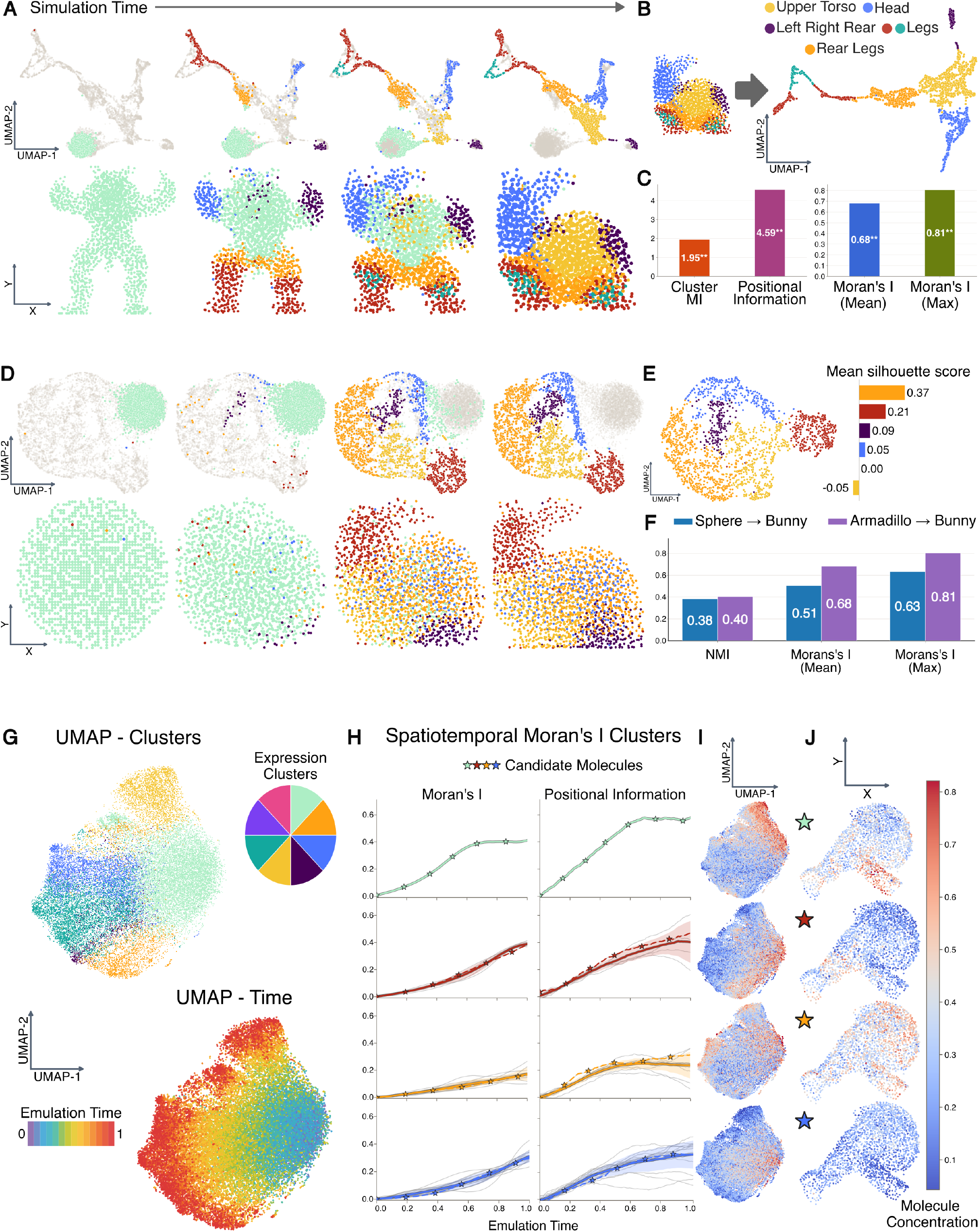
Latent signaling states acquire spatial and temporal organization during learned deformation. **A:** Armadillo-to-Bunny trajectory represented in latent molecular space by uniform manifold approximation and projection (UMAP; top) and in physical space projected onto the *xy* plane (bottom). Cells are colored by Leiden cluster assignment. **B:** Final Bunny morphology colored by latent molecular state cluster, together with the corresponding UMAP. **C:** Positional information and spatial autocorrelation of the final latent states. Left, mutual information between cell position and either Leiden cluster labels (cluster MI) or the continuous latent-state vectors (positional information), reported in bits. Right, global Moran’s *I* for individual latent signals. Asterisks denote significance under permutation testing (1000 repeats, *p <* 0.001). **D:** Sphere-to-Bunny deformation initialized with random latent molecular states and polarities. **E:** Separability of latent molecular states across anatomical regions of the Bunny, quantified by silhouette score. **F:** Comparison of the final latent molecular states learned for sphere-to-Bunny and Armadillo-to-Bunny deformation. The trajectories encode similar positional information but differ in the spatial autocorrelation of their latent signals. **G:** UMAP of latent molecular states sampled across the reconstructed mouse myocardium trajectory, colored by Leiden cluster (top) or trajectory time (bottom) [45]. **H:** Functional clustering of the temporal Moran’s *I* curves for individual latent molecular signals (left) and the positional information associated with each functional cluster over time (right). **I:** UMAP showing representative latent molecular signals from each functional cluster. **J:** Spatial distributions of representative latent molecular signals in the final myocardium, projected onto the *xy* plane.

We next tested whether this organization could emerge when the initial shape provided no anatomical cues. To do so, we learned a deformation from a sphere to the Stanford Bunny, initializing the latent signals and polarities randomly (Fig. 5D). The model broke symmetry and formed distinct signaling domains across the target shape. Regions that moved farther from the initial sphere became more clearly separated in latent-state space, indicating that larger deformations required more distinct local cues (Fig. 5E). The sphere-to-Bunny and Armadillo-to-Bunny trajectories contained similar positional information, but differed in the spatial autocorrelation of their final latent states (Fig. 5F).

We also analyzed the latent signals learned along the mouse myocardium trajectory [45]. A UMAP of states from all frames showed initially similar signals separating into distinct groups over developmental time (Fig. 5G). Because *waxMorph* reconstructs the full trajectory, we tracked the spatial organization of each signal by measuring Moran’s I at every frame. When clustering the resulting curves using functional data analysis (Fig. 5H; *Methods*) [61], we find four clusters of latent signals with distinct spatiotemporal dynamics. Some signals gained spatial organization early, whereas others increased, decreased, or reorganized later in the trajectory (Fig. 5I). These patterns remained visible in the final myocardium, where signals either marked localized regions or varied broadly across the tissue (Fig. 5J). Thus, the learned latent signaling states provide spatial and temporal coordinates that organize tissue deformation.

### Latent *waxMorph* features quantify the information required for shape assembly

In *waxMorph* shape emerges from a latent signaling system which directs locally interacting cells towards a global target morphology. This flexibility can lead to a potential shortcut: if each cell begins with a distinct continuous state, the model could use that state as an identifier and map cells directly to their respective target positions [40]. Random uniform initialization of cell states makes this less likely, but does not rule it out. We therefore evaluated the stability of the learned representations across specimens, model capacities, and initialization schemes.

We evaluated the consistency of the learned representations across biological specimens using the forelimb subset of the mouse limb dataset. The dataset comprises unaffected and *Fgfr2*^+/P253R^ mutant embryos sampled across successive developmental periods. Within each group, we randomly paired same-side forelimb meshes from consecutive periods and learned trajectories for the early-to-mid and mid-to-late transitions (Fig. 6A; *Datasets, Methods*). Target positional information was similar across random pairings within each group, indicating that independently fitted models recover comparable spatial organization in their latent states. In contrast, unaffected and mutant forelimbs differ significantly during the early-to-mid transition (Fig. 6B). This interval coincided with the developmental period in which aberrant *Dusp6* expression was previously reported in *Fgfr2*^+/P253R^ mutants [44].

**Figure 6:**
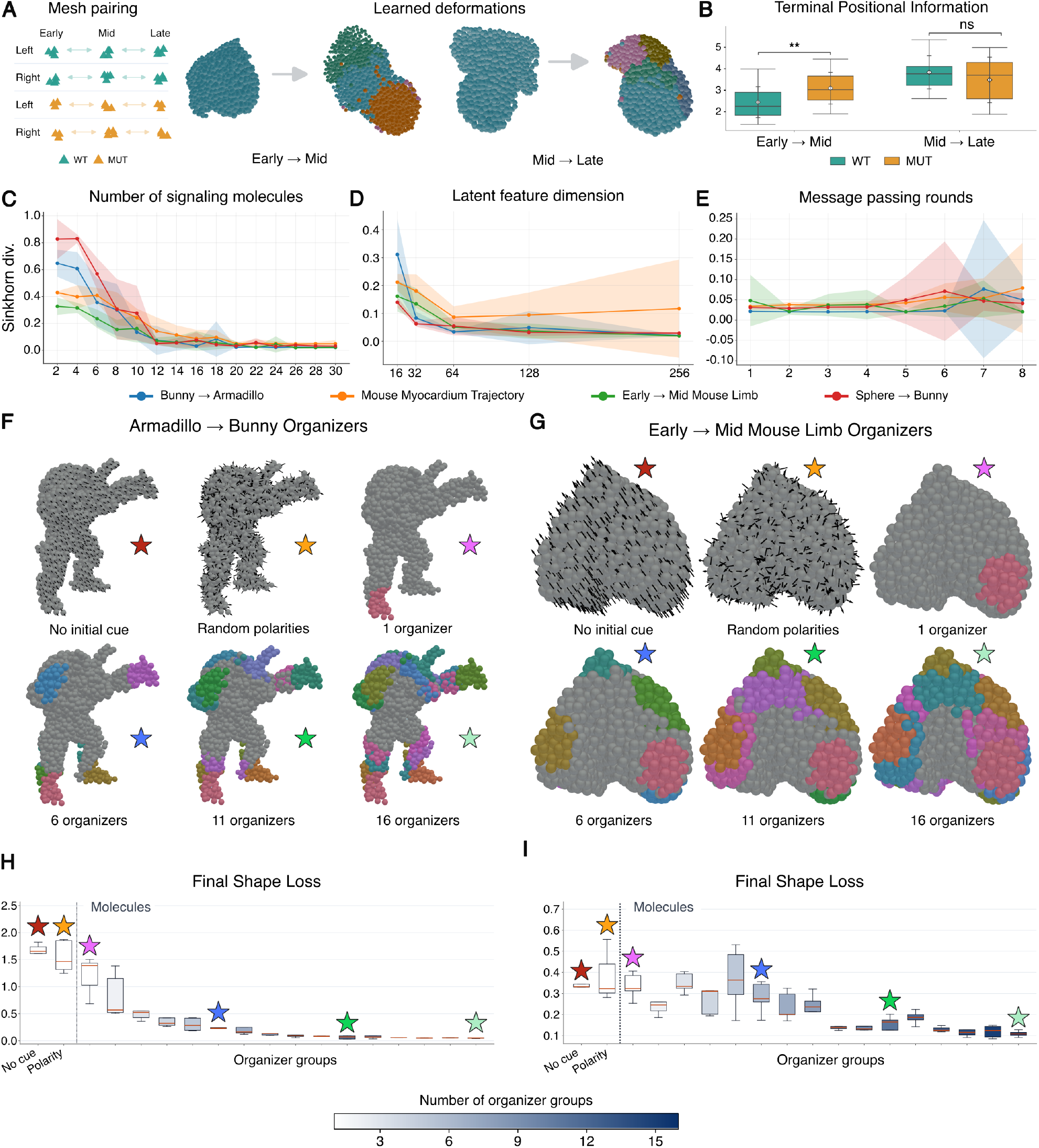
Model capacity and spatial cues reveal requirements for shape assembly. **A:** Analysis of unaffected and mutant mouse forelimbs. Same-side meshes were randomly paired across consecutive developmental periods within each condition (left), and a deformation trajectory was learned for each early-to-mid or mid-to-late pair (right). **B:** Final positional information, measured as the mutual information between cell position and the learned latent signaling state. Independently fitted trajectories show similar values within each group, whereas unaffected and mutant forelimbs differ during the early-to-mid transition. **C–E:** Dependence of final shape reconstruction on model capacity. **C:** Number of latent signaling variables carried by each cell. **D:** Dimension of the latent node and edge representations. **E:** Number of message-passing rounds. **F, G:** Spatial-cue initialization for different deformation tasks with cells colored by organizer. Initial conditions include no spatial cue, random polarity alone, or 1, 6, 11, or 16 localized organizer regions. **H, I:** Final shape loss as a function of organizer number for the Armadillo-to-Bunny (*H* ) and forelimb (*I* ) deformations. The synthetic reconstruction stabilizes with relatively few organizers, whereas forelimb elongation continues to benefit from additional spatial cues. Stars mark the initialization conditions visualized in *F* and *G*.

We next asked how much model capacity was required to reconstruct each deformation. We first varied the number of latent signaling variables carried by each cell. Increasing this number improved shape reconstruction until further variables provided little additional benefit, with the point of saturation differing across tasks (Fig. 6C). We then varied the capacity of the graph neural network that mediates communication between neighboring cells. Increasing the dimension of the latent node and edge features improved reconstruction up to a similar task-dependent limit (Fig. 6E). Increasing the number of message-passing steps initially reduced reconstruction error, but deeper propagation eventually destabilized training (Fig. 6D). This is consistent with oversmoothing in deep graph neural networks, in which repeated aggregation makes neighboring representations increasingly similar. We interpret the capacity beyond which reconstruction no longer improves as an effective intrinsic dimension of the deformation. This provides a model-based measure of assembly complexity: deformations with a higher intrinsic dimension require richer cellular states and communication channels to coordinate shape assembly through local interactions.

We also tested if *waxMorph* could still assemble target morphologies when cell states were not initialized with continuous values. We assigned most cells an identical latent molecular state and designated small, spatially contiguous groups of cells with distinct discrete states. We refer to these localized groups as *organizers*, following developmental signaling regions [10, 40, 62–64]. We also considered conditions without organizers and conditions in which random polarity was the only source of asymmetry (Fig. 6F,G; *Methods*). *waxMorph* still learned deformations across synthetic and real datasets, indicating that reconstruction did not depend on memorizing cell-specific target positions. Introducing organizer regions improves final reconstruction relative to the no-cue and polarity-only conditions. This suggests that a coarse initial spatial pattern can help coordinate intermediate tissue organization during non-uniform deformation. Finally, the number of organizers required differed between tasks. The Armadillo-to-Bunny transformation reached accurate, low-variance reconstructions with relatively few organizers, after which additional cues provided little improvement (Fig. 6H). Forelimb reconstruction was more variable with few or no organizers and continued to improve as additional organizer regions were introduced (Fig. 6I). This shows that forelimb elongation required richer spatial cues than the Armadillo-to-Bunny transformation. In this sense, organizer number provides a model-based measure of how much spatial coordination a form demands.

## Discussion

We developed *waxMorph*, a hybrid physics–deep learning framework in which cells are represented as active agents with molecular features that act under physical constraints. We addressed two complementary questions: what forms emerge from a given set of local rules, and which local rules are sufficient to assemble an observed multicellular morphology or sequence of morphologies?

In the forward direction, *waxMorph* simulates morphodynamic trajectories by coupling molecular dynamics to physical interactions across space and time. Reaction–diffusion systems, mechanical potentials, polarity, growth, and division can all be specified within the same computational framework. Automatic differentiation converts mechanical potentials into force updates, making it easy to specify and explore new physical constraints. We illustrated this flexibility by translating a mechanochemical program previously formulated in a three-dimensional vertex model into a spheroid-based simulator. The resulting epithelial–mesenchymal Turing spheroids recovered elongation, branching, and surface undulation.

In the inverse direction, *waxMorph* learns neighbor-dependent cell state updates while enforcing spatial adjacency, volume exclusion, short-range adhesion, and molecular diffusion. This led to the successful interpolation of sparsely observed tissue morphologies with continuous cellular trajectories. In the developing myocardium, *waxMorph* reconstructed held-out intermediate states more accurately than midpoint interpolation based on optimal transport. The learned latent variables also became organized across space and time, revealing distinct patterns that could be summarized through positional information, spatial autocorrelation, and functional clustering. These variables should not be interpreted as recovered genes. Rather, they describe the internal coordinates the model uses to organize deformation.

The forelimb experiments showed that these representations also captured differences between related biological conditions. Unaffected and *Fgfr2*^+/P253R^ mutant forelimbs differed in terminal positional information during the early-to-mid developmental interval. This coincided with the period in which altered *Dusp6* expression was previously reported. The agreement does not establish a direct correspondence between the learned variables and *Dusp6*, but it suggests that the geometry of the deformation retains information about a known developmental perturbation.

*waxMorph* currently builds shapes by asking whether the cells fill the same three-dimensional region as the target, without requiring each predicted cell to match a particular observed cell. This allows the model to learn from segmented tissue shapes even when cell identities are unknown. When orientation, landmarks, or cell correspondences are available, these can be added to the loss function [40]. The inverse model also keeps the number of cells fixed, which makes it possible to learn tissue deformation and rearrangement while preserving differentiability. Future versions will include differentiable division rules so that cells can be added as the tissue grows.

A practical, yet fundamental advantage of *waxMorph* is that simulation, inference, visualization, and analysis operate on the same evolving tissue representation. Through its integration with Warp and Universal Scene Description [41, 46], trajectories can be rendered, exported, and analyzed without rerunning the underlying computation. The same outputs can be passed to spatial statistics and functional data-analysis tools. Such a shared representation matters because morphogenesis cannot be understood from its final outline alone. Each intermediate geometry bears the trace of the forces, constraints, and local exchanges through which a tissue came to be.

Future work will connect *waxMorph* to single-cell transcriptomics, spatial measurements, live imaging in developing tissues and organoids, and chemical and optogenetic interventions. This will make it possible to ask how cells divide labor, how tissue boundaries emerge and expand, and how molecular programs interact with mechanics to build functional form. Beyond interpolating observed morphologies, *waxMorph* could test competing hypotheses about development. It could ask which cells generate force and which cells respond. It could also reveal how molecular and mechanical memories accumulate and shape cellular responses to changing environments. At its core, morphogenesis is a multiscale, multimodal, multiagent computation. Molecular states, physical interactions, and cellular behaviors are combined through space and time to generate shape. We envision *waxMorph* as a framework for studying this computation with the cell at its core, not only as an object to be modeled and intervened upon, but as the fundamental unit that senses, communicates, acts, and collectively builds the tissue.

## Methods

### waxMorph

*waxMorph* is an ecosystem for modeling cellular interactions and physical constraints that assemble multicellular shape. It is organized into three interoperable components: a forward simulator, a learned emulator, and an interpretability module. The forward simulator integrates prescribed mechanochemical rules; the emulator learns local update rules that assemble a target morphology under prescribed physical constraints; and the interpretability module exports, visualizes, and quantifies the resulting trajectories. The three components operate through the shared cellular representation interface of spheroids and a common set of physical primitives and are implemented in Python with GPU acceleration through NVIDIA Warp [41], PyTorch [42], and JAX [43].

#### Cell state

Each cell is represented as a three-dimensional spheroidal agent. A cell *i* carries a center position **x**_*i*_ ∈ ℝ^3^, a radius *r*_*i*_ ∈ ℝ_+_, a unit polarity vector **p**_*i*_ ∈ ℝ^3^ with ∥**p**_*i*_∥_2_ = 1, and a molecular state. The cell volume is 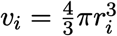, and the concentration **c**_*i,M*_ of a molecule *M* is its abundance divided by *v*_*i*_. The tissue state is the collection 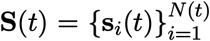 over all *N* (*t*) cells. The molecular state and the auxiliary attributes of a cell are instantiated separately throughout each case study: the forward simulator equips every cell with a two-component activator–inhibitor (*A* –*I* ) pair, an equilibrium radius, and a discrete cell type (*Forward simulator*), whereas the learned emulator equips every cell with a vector of latent signaling molecules and a fixed radius (*Learned emulator*).

#### Dynamic cell neighborhoods

Cell interactions are restricted to spatially adjacent neighbors. The neighborhood graph *G*(*t*) = (*V* (*t*), *E*(*t*)) is reconstructed from cell positions as the tissue deforms, and (*i, j*) ∈ *E*(*t*) denotes that cells *i* and *j* are neighbors at time *t*. The same relation is used to evaluate pairwise mechanical interactions, polarity-dependent potentials, and molecular diffusion. The spatial graph is induced by cell positions rather than by a fixed mesh of cell boundaries, so neighborhoods change as cells move. For a neighbor pair, the center-to-center distance and the unit vector pointing from cell *j* toward cell *i* are

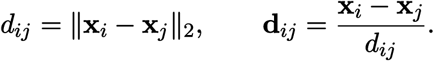

Because spheroidal agents do not define an explicit polygonal contact area, every neighbor pair is assigned the same effective contact area.

#### Hybrid physical primitives

Both the forward simulator and the learned emulator are built from three primitives that act over the neighborhood graph: overdamped mechanics, a reciprocal soft-sphere force computed pairwise for neighboring agents, and graph-based molecular diffusion.

Cell motion is overdamped, so velocity is proportional to the net force, 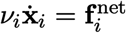, with *ν*_*i*_ an effective friction coefficient [24, 28, 48, 65–69]. The friction coefficient is absorbed into the mechanical parameters and the time scale. Continuous quantities are integrated with the forward Euler method, and polarity vectors are renormalized to unit length after each update.

Volume exclusion and short-range adhesion are imposed by a soft-sphere force between neighbors (*i, j*),

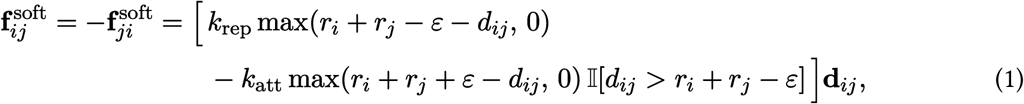

where I[·] is the indicator function, *ε* is the width of the attraction band, and *k*_rep_, *k*_att_ are repulsion and attraction coefficients. The repulsive branch acts for *d*_*ij*_ *< r*_*i*_ + *r*_*j*_ − *ε* and a short-range attractive branch acts over [ *r*_*i*_ + *r*_*j*_ − *ε, r*_*i*_ + *r*_*j*_ + *ε* ].

Molecular transport is evaluated as diffusion over the neighborhood graph. For a concentration field *c*, the discrete graph Laplacian at cell *i* is

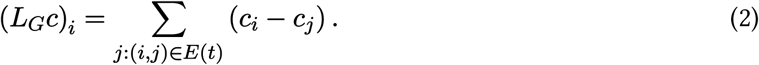

Each case study combined these primitives with additional, case-specific rules.

#### Forward simulator and the learned emulator

In the forward simulator, rules governing mechanical interactions, signaling, growth, and division are prescribed, and the simulator integrates the resulting dynamics. For the learned emulator, the known part of the physics can be prescribed, while the unresolved portion of the local update rule is learned by a graph network-based simulator, so that an initial population assembles a target morphology under physical constraints.

#### Interpretability module

The outputs from the forward simulator and the learned emulator of *waxMorph* can be processed through traditional and functional analysis pipelines. High-quality visualizations and export formats are enabled through *waxMorph*’s rendering module, which is built over OpenUSD [46], OpenGL, PyVista [70] and Matplotlib [71].

### Forward simulator: polarized epithelial–mesenchymal spheroids

As a case study of the forward simulator, we modeled an epithelial–mesenchymal cell aggregate coupled to a two-component activator–inhibitor reaction–diffusion system based on a three-dimensional Turing model [5, 27]. Positions and polarities follow overdamped dynamics, molecular abundances change by reaction and diffusion over the neighborhood graph, and the activator concentration drives mesenchymal growth and division. Mechanical forces were evaluated either from analytically derived gradients (Appendix A) or by automatic differentiation of the corresponding potentials.

#### Cell state

The state of a cell *i* at time *t* was modeled as a vector

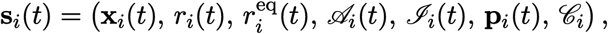

with cell-center position **x**_*i*_ ∈ ℝ^3^, radius *r*_*i*_ ∈ ℝ_+_, equilibrium radius 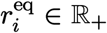, activator and inhibitor abundances *A, I* ∈ ℝ, unit polarity **p** ∈ ℝ^3^, and cell type *C*_*i*_∈ *{*Mes Epi*}*. The number of cells *N* (*t*) changed through division. The volume and molecular concentrations were

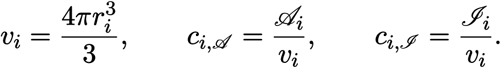

#### Epithelial polarity and thickness potentials

We implemented cell-type-specific potentials to maintain the epithelial monolayer and regulate cell polarity. We defined the epithelial polarity potential as

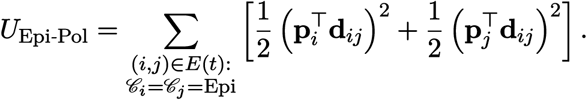

This potential is minimized when neighboring epithelial polarities are perpendicular to their displacement vector, orienting polarity normal to the sheet [28, 50]. To define the epithelial-thickness potential, we first aligned the signs of neighboring polarities for each epithelial pair (*i, j*),

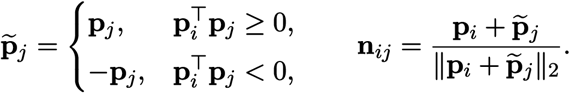

The resulting epithelial-thickness potential,

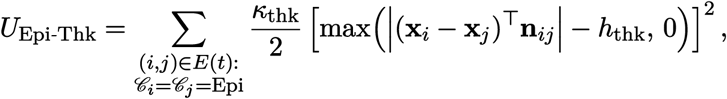

penalizes displacement of neighboring epithelial cells along the local sheet normal, thereby reducing cell stacking. We implemented the position-dependent potential and its corresponding force as

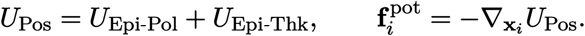

#### Polarity dynamics

Mesenchymal polarity was regulated by two potentials. The first aligned the polarity axes of neighboring mesenchymal cells,

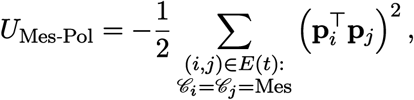

without distinguishing orientation as the inner product is squared. The second aligned the polarity of a mesenchymal cell toward neighbors of higher activator concentration,

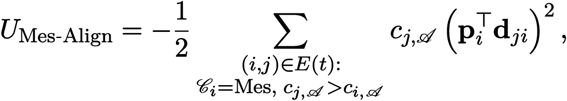

where the higher-activator neighbor *j* could be mesenchymal or epithelial. The resulting polarity potential and dynamics are

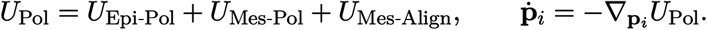

#### Mechanical updates

Position updates combined the cell-type-specific potential force with the shared soft-sphere force (Eq. 1),

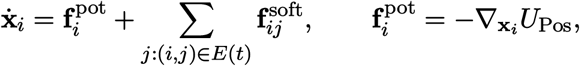

with soft-sphere coefficient *k*_rep_ for all pairs and *k*_att_ set per cell-type pair (Sup. Tab. 2). Positions and polarities were integrated by forward Euler with step Δ*t*_mech_,

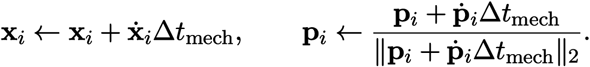

Gradients of the mechanical potentials were evaluated either from the analytical expressions in Appendix A or by automatic differentiation in Warp [41]; in the latter, position and polarity updates were obtained directly from the scalar potentials *U*_Pos_, *U*_Pol_ using wp.grad.

#### Activator–inhibitor reaction–diffusion

Chemical signaling used a two-component activator–inhibitor system [5, 27, 49]. For cell *i*, abundances evolved as

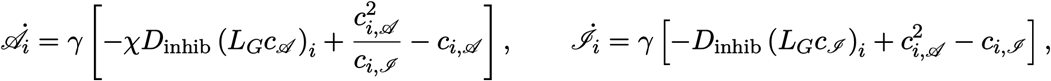

where the diffusion term uses the shared graph Laplacian (Eq. 2), *γ* is the reaction time scale, *D*_inhib_ is the inhibitor diffusivity, and 0 ≤ *χ* ≤ 1 is the activator diffusivity relative to that of the inhibitor (the spatial characteristic). Abundances were integrated by forward Euler with step Δ*t*_chem_,

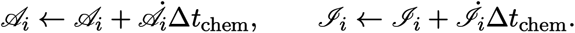

#### Activator-dependent growth

Mesenchymal growth was controlled by the intracellular activator concentration. The equilibrium radius followed a Hill function,

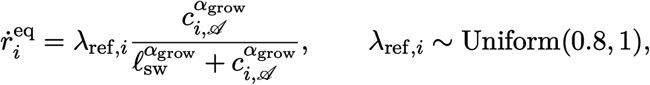

with switch concentration *ℓ*_sw_ and Hill exponent *α*_grow_. The physical radius relaxed toward the equilibrium radius,

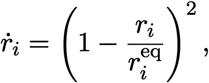

and both were integrated with step Δ*t*_grow_,

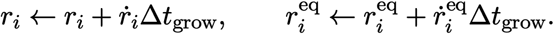

#### Division

We set the probability that a mesenchymal cell *i* divided during a step to

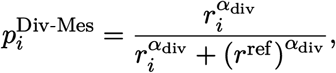

with reference radius *r*^ref^ and exponent *α*_div_, producing a steep increase in division probability as the radius approached *r*_max_. Epithelial cells did not undergo activator-dependent growth; an epithelial cell was instead marked for division when it had at least one mesenchymal neighbor and fewer than six epithelial neighbors, adding epithelial cells as the enclosed tissue expanded while maintaining the monolayer. Cells marked for division were processed in parallel. A mesenchymal parent was replaced by two daughters of the same type and polarity, with molecules divided equally,

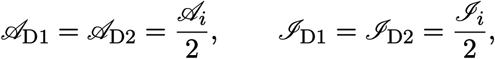

and radii initialized to conserve volume,

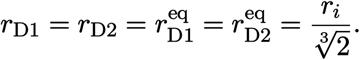

For epithelial divisions, the parent radius was copied to both daughters, 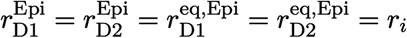, allowing the epithelial layer to expand as the enclosed mesenchyme grew [28]. The simulator supports division axes parallel or perpendicular to the parent polarity. For a polarity-perpendicular division, a reference vector **u**_ref_ not parallel to **p**_*i*_ was chosen and

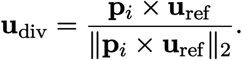

Daughter centers were placed on opposite sides of the parent center,

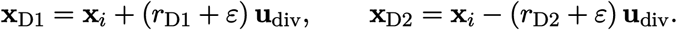

#### Initial conditions

Simulations were initialized with *N*_0_ cells (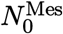 mesenchymal, 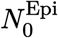 epithelial; see Sup. Tab. 2). Epithelial cells were arranged as a monolayer on a Fibonacci lattice over a sphere of radius *R*_shell_ and were initialized with no activator or inhibitor. Mesenchymal cells were sampled uniformly inside a ball of radius *R*_inner_. Their initial activator and inhibitor abundances followed a 3D vertex model with an activator-to-inhibitor ratio of 5.0 [27]. All cells were initialized with radii 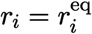 and polarities 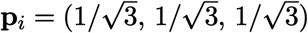. Before growth, the mechanochemical dynamics were run for 60,000 steps.

#### Automatic-differentiation comparison

Simulations in which mechanical updates were obtained from analytically derived gradients were compared with simulations in which the same updates were obtained by automatic differentiation of the mechanical potential. Two dynamical regimes were evaluated: an elongation–branching regime (*χ* = 0.005, *γ* = 0.001) and a surface-undulation regime (*χ* = 0.005, *γ* = 0.01). For each regime and initial condition, two trajectories were generated with analytically derived updates and one with automatic differentiation, using distinct random seeds but identical initial configuration and parameters. The two analytical trajectories estimate the variation caused by nondeterministic GPU execution, including floating-point atomic reductions and random memory-access order; the difference between an analytical and an automatic-differentiation trajectory therefore includes both this baseline variation and any difference introduced by the gradient-evaluation method. *N*_rep_ = 20 independent experiments were generated per regime, and each tissue was simulated until it contained 80,000 cells.

#### Shape alignment

Morphologies were aligned over the special Euclidean group SE(3). Each morphology was first centered at its centroid to remove translation. Rotational differences were then removed by optimizing a three-dimensional rotation. Following [72], for **z** = (*z*_1_, *z*_2_, *z*_3_)^⊤^, the skew-symmetric matrix

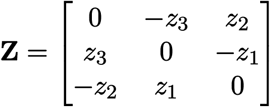

defines the Cayley rotation

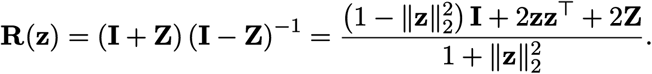

The rotation parameters were estimated by minimizing a differentiable point-set loss between the centered source and rotated target,

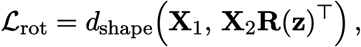

with *d*_shape_ the debiased Sinkhorn divergence from GeomLoss [73]. The loss was optimized with Adam at learning rate 0.05.

#### Shape-matching distance

We compared aligned morphologies using a symmetric, normalized Chamfer distance (NCD). For two point sets 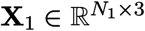 and 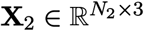,

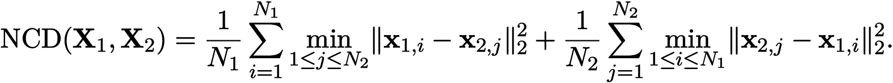

The NCD was reported for terminal morphologies and at intermediate states sampled after each increase of 1,000 cells.

### Inverse emulator: learned emulation of morphogenesis

As a case study of the inverse mode, biophysically constrained tissue deformations were learned with a graph-network simulator [35, 74, 75]. The known soft-sphere mechanics (Eq. 1) and graph-based diffusion (Eq. 2) were prescribed as biophysical constraints, and neighbor-dependent updates to position, polarity, and latent molecular state were learned so that an initial population assembled target volumes.

#### Cell state

The emulated system was represented over discrete steps *t* = 0, …, *T* . The state at time *t* concatenated a fixed number *N* of agents, 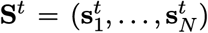, and a rollout was the sequence **S** = (**S**^0^, …, **S**^*T*^ ). Each agent state was

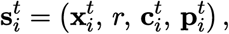

with position **x**_*i*_ ∈ ℝ^3^, constant uniform radius *r* (so 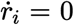), latent signaling molecule concentrations 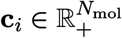, and unit polarity **p**_*i*_ ∈ ℝ^3^. The number of agents was fixed throughout rollouts.

#### System-state initialization

Initial states were generated from a mesh describing the target volume by using *waxMorph*’s data module, sampling spheroid agents representing the mesh volume for all datasets (*Datasets*). Meshes were loaded, repaired, reoriented, and normalized with trimesh [76]. For a requested particle count *N* and mesh volume *V*, a uniform particle radius was set as

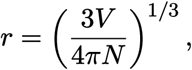

with exclusion distance *d*_min_ = 1.2 *r*. The mesh was voxelized at pitch 0.4 *d*_min_; when too few voxels resulted, binary dilation, hole filling, and erosion were applied with SciPy [77]. Candidate voxels were thinned in random order (with an optional fixed seed) toward an average spacing of *d*_min_, and a candidate center **x**_*i*_ was accepted if and only if no accepted center **x**_*j*_ satisfied ∥**x**_*i*_ **x**_*j*_∥_2_ *d*_min_, similar to dart-throwing for Poisson-disk sampling [54–56]. Latent concentrations and polarities were initialized randomly,

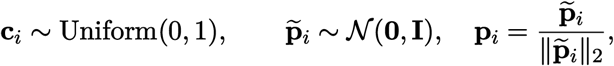

unless a structured initialization was used (see *Spatial-cue initialization*).

#### Deformation task

The objective was to deform the system so that the agent centers 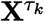 matched the volume described by the target samples 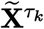 at *K* specified time points along the trajectory, encoded as goal tuples 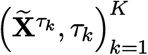 with 1 ≤ *K* ≤ *T* . Targets were sampled from meshes with the same data module. The shape loss summed a distributional distance over goal time points,

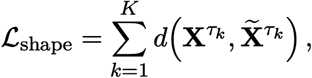

reducing to 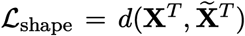 for a single terminal target. Because the rows of 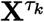 and 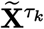 are unordered, *d*(·,·) was a distributional distance between point sets. Specifically, *waxMorph* provides access to Chamfer distance [78] and GeomLoss [73] implementations of maximum mean discrepancy [79], Hausdorff divergence [80], and debiased Sinkhorn divergence [73]. Experiments used the default sinkhorn divergence, a fast approximation of the 2-Wasserstein distance [73, 81]. With empirical measures

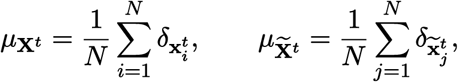

the 2-Wasserstein distance was

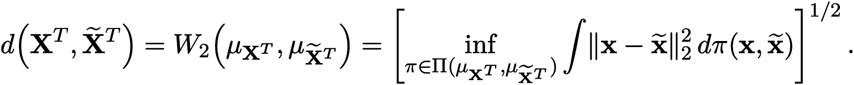

A regularizer *λ* penalized large position changes between consecutive frames, encouraging smoother trajectories with minimal movement:

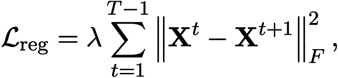

where ∥·∥_*F*_ is the Frobenius norm.

#### Graph-network emulator

Each emulation step maps **S**^*t*^ to **S**^*t*+1^ through a graph neural network architecture [74, 75] that learns neighbor-dependent updates, followed by application of the prescribed biophysical constraints. The network differs from mesh-based simulators in three respects: node features carry the internal signaling state **c**_*i*_ while edge features carry mechanical quantities; the graph is induced by spatial adjacency rather than by a fixed mesh [74] or a fully connected graph [40]; and learned updates are applied together with prescribed biophysical behavior, namely soft-sphere volume exclusion with short-range adhesion (Eq. 1) and molecular diffusion (Eq. 2). One step proceeds through graph construction, latent encoding, message passing, decoding to Euler updates, and enforcing biophysical constraints. At each step the summand of *L*_reg_ and, when a goal time point coincides with *t* + 1, the summand of *L*_shape_ were accumulated.

#### Spatial adjacency graph

Given a state **S**^*t*^, the graph *G*^*t*^ = (**V**^*t*^, **E**^*t*^) was constructed with one node per agent and node features equal to the signaling concentrations, 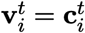. An edge 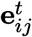 connected agents *i* and *j* if and only if

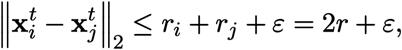

and its features concatenated the center distance 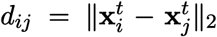 and the polarity angle *θ*_*ij*_ = arccos 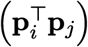. Reconstructing the graph at each step, combined with spatial hashing, yields *O*(*N* ), rather than *O*(*N* ^2^) computational cost, allowing *waxMorph* to model volumes with thousands of agents.

#### Latent encoding

Node and edge features were embedded by multilayer perceptrons *f*_*φ*_ and *f*_*ρ*_ [74, 75],

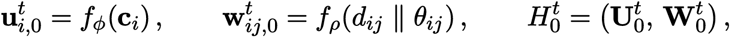

where (· ∥ ·) denotes concatenation. The latent graph 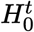 and all subsequent latent graphs share the topology of *G*^*t*^.

#### Message passing

The latent graph was updated by *M* graph-network blocks [35], each preserving topology and updating only the latent node and edge attributes. Because latent messages are directional, *i* → *j* and *j* → *i* were computed separately. At step *m*, a message was assembled from the sender, receiver, and edge embeddings,

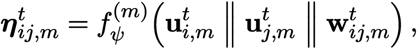

the edge embedding was updated residually,

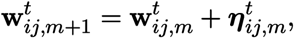

incoming messages were aggregated,

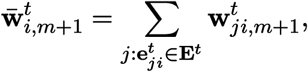

and the node embedding was updated residually through 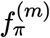,

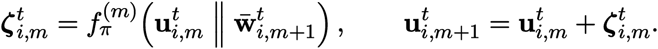

After *M* blocks, the final latent graph was 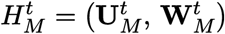; the number of message-passing steps *M* is a hyperparameter (Sup. Tab. 2).

#### State decoding and integration

The final node embeddings, which summarize the information from each agent’s neighborhood, were decoded into state updates using multilayer perceptrons *f*_*ω*_, *f*_*µ*_, *f*_*ν*_,

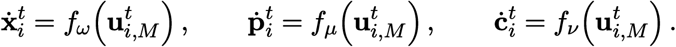

The resulting updates were applied using forward Euler integration with step Δ*t*,

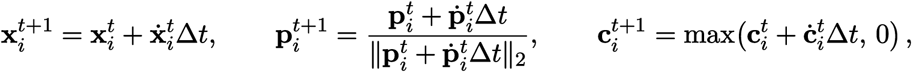

where polarity was renormalized and concentrations were clamped to nonnegative values.

#### Enforcing biophysical constraints

To obtain the full update, the learned updates were followed by enforcing biophysical constraints. The soft-sphere force (Eq. 1) with *r*_*i*_ = *r*_*j*_ = *r* was applied to the provisional positions,

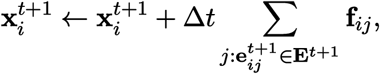

where the adjacency graph was rebuilt from the provisional positions to obtain **E**^*t*+1^. Each latent molecule *M* diffused independently through the shared graph Laplacian (Eq. 2),

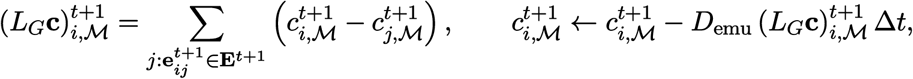

with diffusion coefficient *D*_emu_. The constraints were applied *n*_substeps_ times per learned update on a faster time scale to keep the trajectory biophysically coherent, similar to using different time steps across equations during forward integration of simulators [27]. The next state was 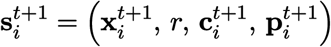. Additional constraints can be added through torch.autograd.Function classes, supported internally by Warp [41].

#### Optimization

The total trajectory loss was calculated as *L* = *L*_shape_+*L*_reg_. The network parameters,

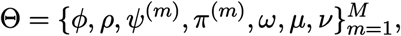

were updated using the AdamW [82, 83] optimizer. Gradient checkpointing [84, 85] was used by default to reduce memory at the cost of recomputation during backpropagation.

#### Spatial-cue initialization

For the constrained-initialization regime, the random concentration initialization was replaced by a controlled spatial asymmetry inspired by localized organizer regions [10, 40, 62–64]. With *K* organizer regions, polarities were random unit vectors and concentrations were initialized uniformly at 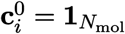; then for each organizer group *A*_*k*_, *k* = 1, …, *K*, of *N*_org_ cells,

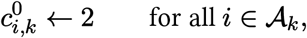

leaving all other entries at one. Organizer groups were selected from the outer shell of the source point cloud. The cloud was centered,

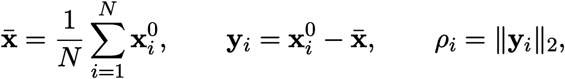

and for shell fraction *f*, the candidate shell was the radial quantile

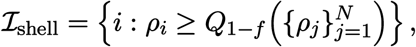

expanded to the *KN*_org_ cells of largest *ρ*_*i*_ if it held fewer than *KN*_org_ candidates. Organizer directions were placed as a nested, well-separated set on Σ^2^: the first eight were the normalized cube vertices 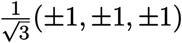, and additional directions were chosen from a deterministic Fibonacci-sphere candidate set [86] by greedily minimizing the maximum dot product with the already selected directions. For each direction **d**_*k*_, the pole was the extreme opposing point,

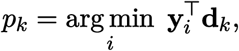

and *A*_*k*_ comprised the *N*_org_ unused shell candidates closest to (and including) *p*_*k*_. Evaluated configurations used *K* ∈ *{*1, …, 16*}* organizers together with two *K* = 0 baselines (uniform and random polarities).

#### Interpolation benchmarks

For endpoint interpolation, *waxMorph* was compared with DiffeoMorph [40], an SE(3)-equivariant point-cloud method. The authors’ implementation was used with the same number of training epochs (2,000) and the same volumetric sample of 2,000 agents as *waxMorph*, with the USE_ATTENTION and USE_ANGLE flags enabled and the remaining hyperparameters matched where possible. For densely sampled trajectories in the leave-one-out setting, *waxMorph* was compared with WaddingtonOT (WOT) [57], which learns optimal-transport couplings between observed frames. Treating coupled frames as animation keyframes, held-out frames were interpolated as the element-wise midpoint of coupled positions **x**^*t*^ and **x**^*t*+2^,

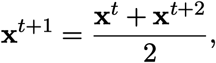

with couplings from compute_transport_matrix using optimal_transport_duality_gap at default settings.

#### Implementation

The emulator is implemented in both JAX [43] and PyTorch [42]; the PyTorch path is the default option for *waxMorph*, as Warp’s automatic differentiation integrates through torch.autograd.Function, while the JAX path requires compile-time array sizes and an upper bound on the number of edges and therefore requires more memory. Unless stated otherwise, network width, latent dimension, trajectory length, number of latent molecules, learning rate, and epochs are as listed in Sup. Tab. 2. All networks used the sigmoid-weighted linear unit (SiLU) activation [87].

### Rendering module and interpretability analysis

#### Supported rendering backends

*waxMorph* supports high-quality output formats for 3D rendering directly to videos (via OpenGL), or exporting to USD files [46, 47] which can then be rendered or viewed in third-party software. Through its render module, *waxMorph* provides multiple bridges for utilizing static backends for 2D and 3D renders through Matplotlib [71] and PyVista [70], or dynamic backends for videos through Warp. The repository includes examples of using these backends, both for simulated and learned trajectories.

#### Traditional analysis of emulated trajectories

The clustering analyses for signaling molecules on synthetic and real examples were conducted through scanpy [88] to embed the data into UMAPs [58] and perform Leiden clustering [89].

For sampled trajectories in the synthetic and real deformation examples, UMAP was used with *n*_neighbors_ = 30, metric = cosine, min_dist = 0.15. Learned latent molecular signaling concentrations were normalized using the StandardScaler provided through scikit-learn [90]. The resolution parameter for Leiden was set to 0.3.

For the interpretability analysis using the developing myocardium data [45], we visualized the resulting latent information via UMAP with settings: *n*_neighbors_ = 75, metric = euclidean, min_dist = 0.15. We performed Leiden clustering with a 0.6 resolution parameter.

To quantify positional information (PI), defined as the mutual information (MI) between molecule concentrations and positions, we use two approaches:

- With clustering, we calculated the mutual information between signaling molecule concentration clusters *y* and positions **x**_*i*_. Cluster labels were obtained using the Leiden algorithm [89].
- Without clustering, we calculated the mutual information between concentrations **c**_*i*_ and positions **x**_*i*_ directly.

In both cases, the mutual information is quantified using the Kraskov-Stögbauer-Grassberger (KSG) estimator implemented using the infomeasure [59] package.

We used Global Moran’s I to measure spatial autocorrelation [60, 91]. Let 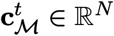 denote the concentrations of a single molecule *M* across all agents at time *t*,

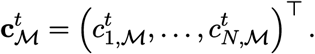

Let **A**^*t*^ ∈ ℝ^*N ×N*^ denote the row-normalized *K*-nearest-neighbor spatial weights matrix at time *t*, with *K* = 75 and 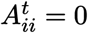. Then Global Moran’s I for molecule *M* at time *t* is

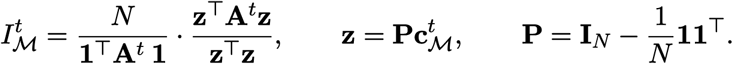

#### Functional analysis of learned heart trajectory

The densely sampled emulator trajectories obtained from *waxMorph* can be used to conduct functional analyses, such as spatiotemporal clustering [92]. We performed this analysis in the following steps:

1. We calculated Global Moran’s I [60] and approximate positional information (PI) [8] for each molecule and timepoint along the trajectory. This yields an empirical record of Global Moran’s I as a function of trajectory time.
2. We used scikit-fda [61], particularly the K-Means algorithm for functional data (KMeans), for different values of the hyperparameter *K* ∈ [2, 6]. We picked the final *K* yielding the highest silhouette score (i.e., highest separability) across curves, calculated through silhouette_score in scikit-learn.

## Datasets

All data volumes were sampled with the data module ( *System-state initialization*) using *N* = 2,000 agents.

### Synthetic meshes

The sphere mesh was generated using the trimesh.creation.icosphere function with subdivisions = 4 and radius = 1. The Stanford Bunny [53] and Armadillo [52] meshes were downloaded from the Stanford 3D Scanning Repository. All synthetic meshes are available in the main repository.

### Wild-type and mutant mouse forelimb [44]

Data for meshes, including information on limb type (WT, mutant), side (right, left), and time points were downloaded through [the original publication repository hosted over Dryad](https://datadryad.org/dataset/doi:10.5061/dryad.8h646s0).

### Developing mouse myocardium [45]

Video frames and 3D meshes for each time point of the heart deformation data were downloaded from [the original publication repository](https://data.mendeley.com/datasets/t828xhg66k/1).

## Code availability

*waxMorph* is available and maintained by the Morpho Lab at its [GitHub](https://github.com/Computational-Morphogenomics-Group/waxmorph) repository. Examples for running forward simulations and the inverse learning pipeline are provided through guided notebooks.

## Acknowledgements

We thank members of the Dumitrascu group (Morpho Lab) for their comments and suggestions. We are grateful to Sebastian Salazar for his input on the design of the physics engine, and to Onur Beker, Kim Stachenfeld, Ryan P. Adams for early feedback on ideas related to this work. We thank Karen Kasza, Kat Hadjantonakis, Jakub Sedzinski for continued conversations about morphogenesis; Noah O. ládélé for feedback regarding the writing of this manuscript. This work was supported by NIH grant 1R35GM157082-01. B.D. was also supported by a CIFAR MacMillan Multiscale Human Fellowship.

## Appendix A Derivation of the mechanical potential

To compare with automatic differentiation, we derive the closed-form gradient updates for the mechanics potentials in the forward simulator. Specifically, we are interested in the quantities:

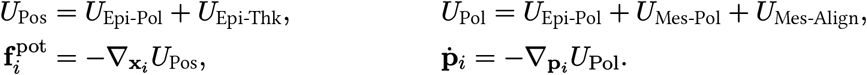

This requires calculating the following quantities:

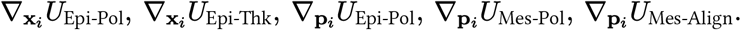

### A.1 Position updates

#### A.1.1 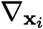 *U*Epi-Pol

We first derive the contribution from one epithelial pair (*i, j*), and then sum over all epithelial neighbors of cell *i*. Recall that:

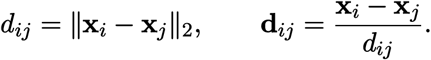

When differentiating with respect to **x**_*i*_, the polarities are treated as fixed. For any polarity vector **p**:

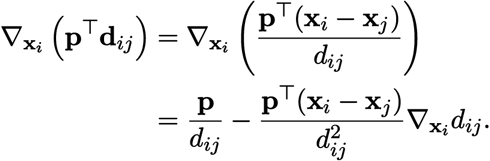

Since 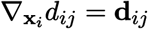 and **p**^⊤^(**x**_*i*_ − **x**_*j*_) = *d*_*ij*_**p**^⊤^**d**_*ij*_, this becomes:

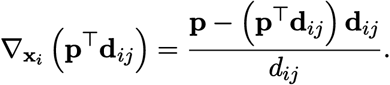

Now applying the chain rule to one pairwise epithelial-polarity term gives:

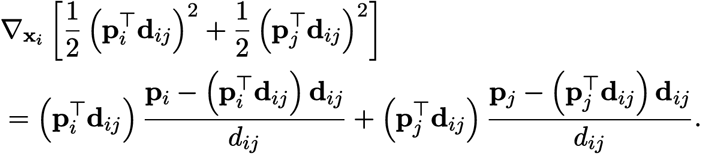

Therefore, the full gradient is:

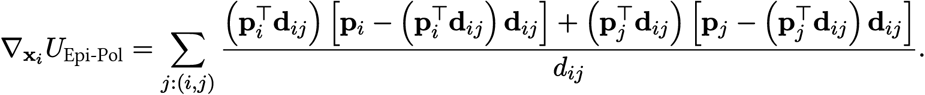

#### A.1.2 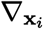 *U***Epi-Thk**

The epithelial thickness potential *U*_Epi-Thk_ is defined as:

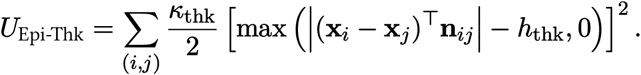

When differentiating with respect to **x**_*i*_, the interpolated polarity **n**_*ij*_ is treated as fixed. For one epithelial pair, the derivative of the scalar projection is:

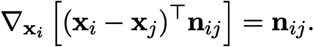

The absolute value contributes a sign:

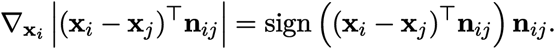

When |(**x**_*i*_ − **x**_*j*_)^⊤^**n**_*ij*_ |≤ *h*_thk_, the maximum term is zero and this pair contributes no gradient. Otherwise, applying the chain rule gives:

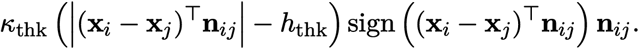

Equivalently, writing the active and inactive cases together:

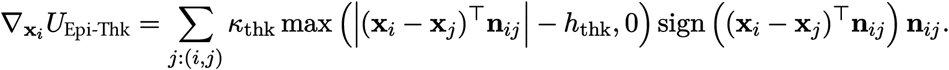

At the non-differentiable point where the maximum is exactly zero, we use the zero subgradient.

### A.2 Polarity updates

#### A.2.1 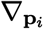 *U*Epi-Pol

For one epithelial neighbor pair (*i, j*), the contribution to *U*_Epi-Pol_ is:

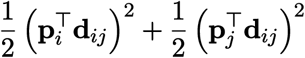

When differentiating with respect to **p**_*i*_, the positions are treated as fixed. Therefore:

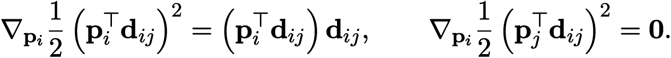

Summing over epithelial neighbors gives:

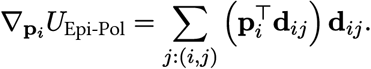

#### A.2.2 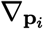 *U***Mes-Pol**

For one mesenchymal neighbor pair (*i, j*), the pairwise contribution is:

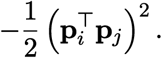

Applying the chain rule:

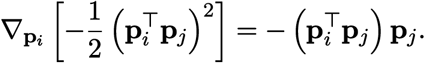

Summing over mesenchymal neighbors gives:

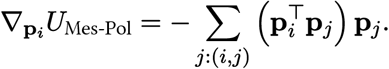

#### A.2.3 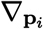 *U***Mes-Align**

For one active neighbor pair (*i, j*) with *I*_*i*_ = Mes and *c*_*j,A*_ *> c*_*i,A*_, the pairwise contribution is:

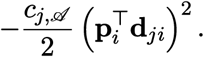

When differentiating with respect to **p**_*i*_, the positions and activator concentrations are treated as fixed. Applying the chain rule:

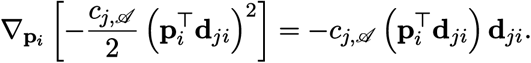

Summing over active neighbors gives:

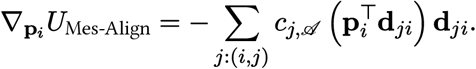

## Appendix B Supplementary materials

**Supplementary Table 1:** Comparison of *waxMorph* with other 3D biophysical modeling frameworks simulating and emulating biophysics. N/A denotes features out of the scope of given methods.

|  | Simulators<br>[27–29, 93] | DiffeoMorph [40] | jax-morph [39] | *waxMorph* (Ours) |
| --- | --- | --- | --- | --- |
| Tissue Model | Spheroids / V. M. | Point Clouds | 2D Spheroids<sup>1</sup> | 3D Spheroids |
| Forward Simulation | ✓ | N/A | N/A | ✓ |
| Simulator Autodiff Support | ✗ | N/A | N/A | ✓ |
| Learned Emulation | N/A | ✓ | ✓ | ✓ |
| Emulating biophysical constraints | N/A | $\sim^2$ | ✓ | ✓ |
| Communication graph | N/A | fully-connected,<br>attention-weighted<sup>3</sup> | spatial | spatial<sup>4</sup> |
| Emulation task | N/A | SE(3)-equiv. shape<br>matching | Various (e.g.<br>elongation) | Density-based<br>shape matching |
| GPU-accelerated rendering | ✓ | ✗ | ✗ | ✓ |
| GPU Support | CUDA/C++ | JAX | JAX | Warp+PyTorch/JAX |

<sup>1</sup> Does not consider polarized spheroids [39].
<sup>2</sup> Does not explicitly prescribe physics, but instead learns a force model and signaling network from scratch [40].
<sup>3</sup> $\mathcal{O}(n^2)$ computational cost with number of agents.
<sup>4</sup> With spatial hashing, achieves roughly $\mathcal{O}(n)$ computational cost with number of agents.

**Supplementary Table 2:** Model and simulation parameters for the forward simulator and the learned emulator. Analysis settings for the interpretability and rendering module are given in the corresponding subsection.

|  | Symbol | Description | Value |
| --- | --- | --- | --- |
| Forward simulator | $k_{\text{rep}}$ | Soft-sphere repulsion coefficient | 3.0 |
| | $k_{\text{att}}$ | Soft-sphere attraction coefficient (Epi–Epi; other pairs) | 1.5; 0.15 |
| | $\varepsilon$ | Attraction-band width | 0.01 |
| | $\kappa_{\text{thk}}$ | Epithelial-thickness stiffness | 6 |
| | $h_{\text{thk}}$ | Epithelial-thickness offset | 0.08 |
| | $\Delta t_{\text{mech}}$ | Mechanical Euler step size | 0.05 |
| | $D_{\text{inhib}}$ | Inhibitor diffusivity | 10 |
| | $\Delta t_{\text{chem}}$ | Reaction–diffusion Euler step size | 0.05 |
| | $\chi$ | Spatial characteristic (relative activator diffusivity) | 0.005 |
| | $\gamma$ | Reaction rate (elongation–branching; undulation) | 0.001; 0.01 |
| | $\ell_{\text{sw}}$ | Growth Hill switch concentration | 0.5 |
| | $\alpha_{\text{grow}}$ | Growth Hill exponent | 10 |
| | $\lambda_{\text{ref},i}$ | Growth reference rate | $\sim \text{Uniform}(0.8, 1)$ |
| | $\Delta t_{\text{grow}}$ | Growth Euler step size | 0.005 |
| | $r^{\text{ref}}$ | Division reference radius | 1.0 |
| | $\alpha_{\text{div}}$ | Division Hill exponent | 20 |
| | $r_{\text{max}}$ | Maximum radius | 1.4 |
| | $N_0 (N_0^{\text{Mes}}, N_0^{\text{Epi}})$ | Initial cell count (mesenchymal, epithelial) | 100 (14, 86) |
| | $R_{\text{shell}}$ | Epithelial shell radius | 3 |
| | $R_{\text{inner}}$ | Mesenchyme ball radius | 1.8 |
| | $r_i(0) = r_i^{\text{eq}}(0)$ | Initial radius | 0.6 |
| | $\mathcal{A}_i(0), \mathcal{I}_i(0)$ | Initial mesenchyme abundances | 0.8, 0.16 |
|  | – | Pre-pattern steps before growth | 50,000 |
| | $N_{\text{rep}}$ | Replicates per regime | 20 |
|  | – | Cells at termination | 80,000 |
|  | – | Adam learning rate (shape alignment) | 0.05 |
| Learned emulator | $r$ | Uniform particle radius | $(3V/4\pi N)^{1/3}$ |
| | $d_{\text{min}}$ | Sampling exclusion distance | $1.2 r$ |
| | – | Voxelization pitch | $0.4 d_{\text{min}}$ |
| | $N$ | Agents per volume | 2,000 |
| | $\mathbf{c}_i(0)$ | Initial latent concentrations (non-organizer) | $\sim \text{Uniform}(0, 1)$ |
| | $N_{\text{mol}}$ | Latent signaling molecules | 32 |
| | $M$ | Message-passing steps | 5 |
| | – | Hidden layers × width per network | $2 \times 256$ |
|  | – | Latent node/edge dimension | 256 |
| | $\Delta t$ | Emulation Euler step size | 0.01 |
| | $D_{\text{emu}}$ | Diffusion coefficient | 0.1 |
| | $n_{\text{substeps}}$ | Constraint substeps per learned update | 5 |
| | $\lambda$ | Trajectory regularization strength | 0.001 |
| | $T$ | Trajectory length | 100 |
|  | – | AdamW learning rate | 0.001 |
|  | – | Training epochs | 2,000 |
| | $f$ | Organizer shell fraction | 0.35 |
| | $N_{\text{org}}$ | Cells per organizer | 50 |
| | $K$ | Organizer regions | $\{1, \dots, 16\}$ |
|  | – | Concentration at organizer cells (vs. background) | 2 (vs. 1) |

**Supplementary Figure S1:**
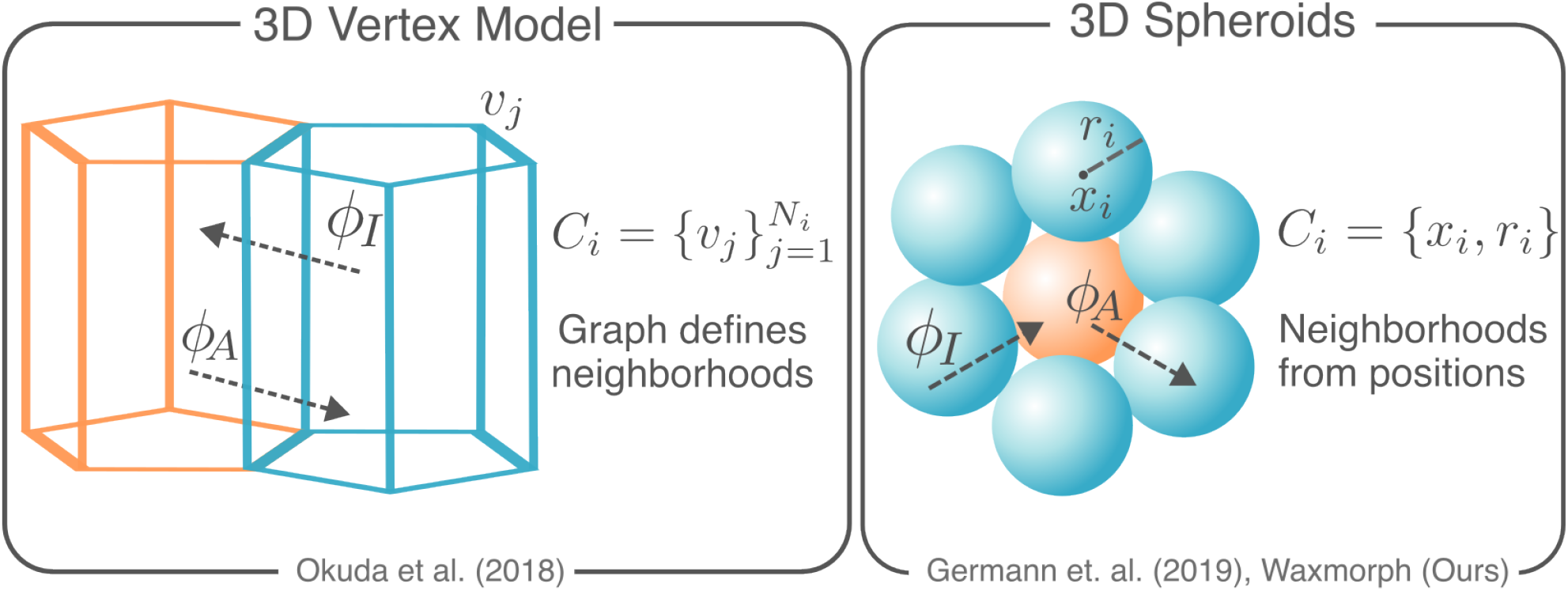
Vertex models and spheroids. In the 3D vertex model, cells establish neighborhoods based on boundaries designated by a graph (left). On the other hand, for 3D spheroids, cell neighborhoods are induced by positional proximity (right).

**Supplementary Figure S2:**
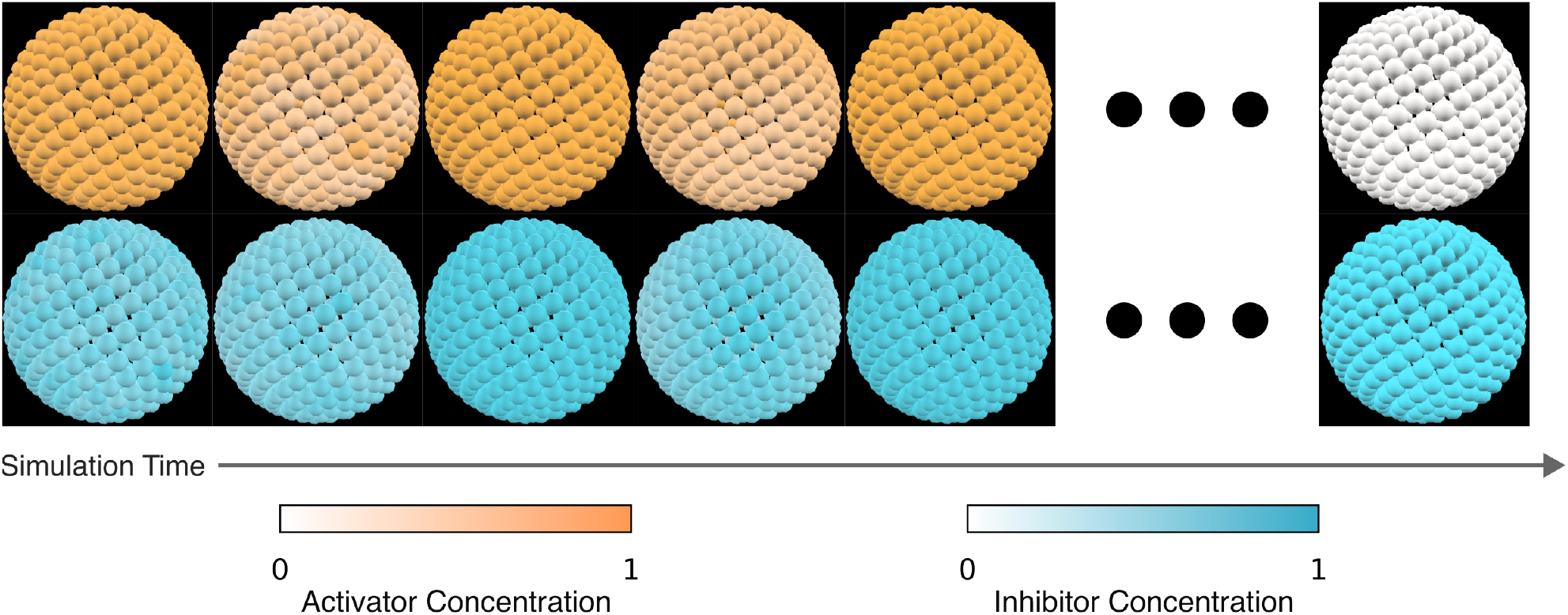
Effect of too high spatial characteristic. Relative activator (top) and inhibitor (bottom) concentrations across cells for different timepoints of the simulation. The monolayer pulses multiple times before converging to a state where patterning is lost.

**Supplementary Figure S3:**
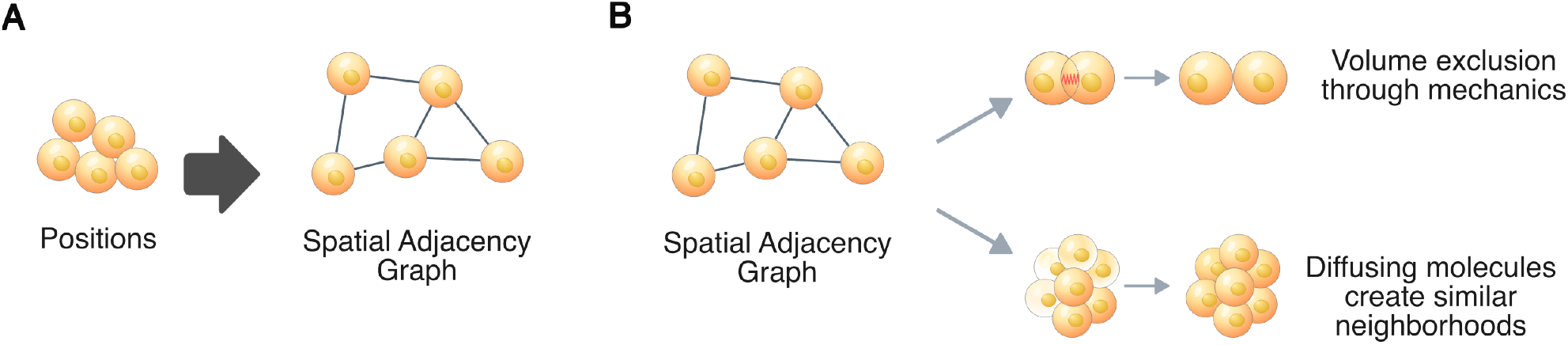
Physical constraints for the emulation experiments. **A**. The state is encoded as a spatial adjacency graph, constructed from positions and state variables. **B, top**. Volume exclusion is attained through using a force term that pushes cells apart. **B, bottom**. Diffusion of molecules drive similar neighborhoods, increasing patterns with high spatial autocorrelation.

**Supplementary Figure S4:**
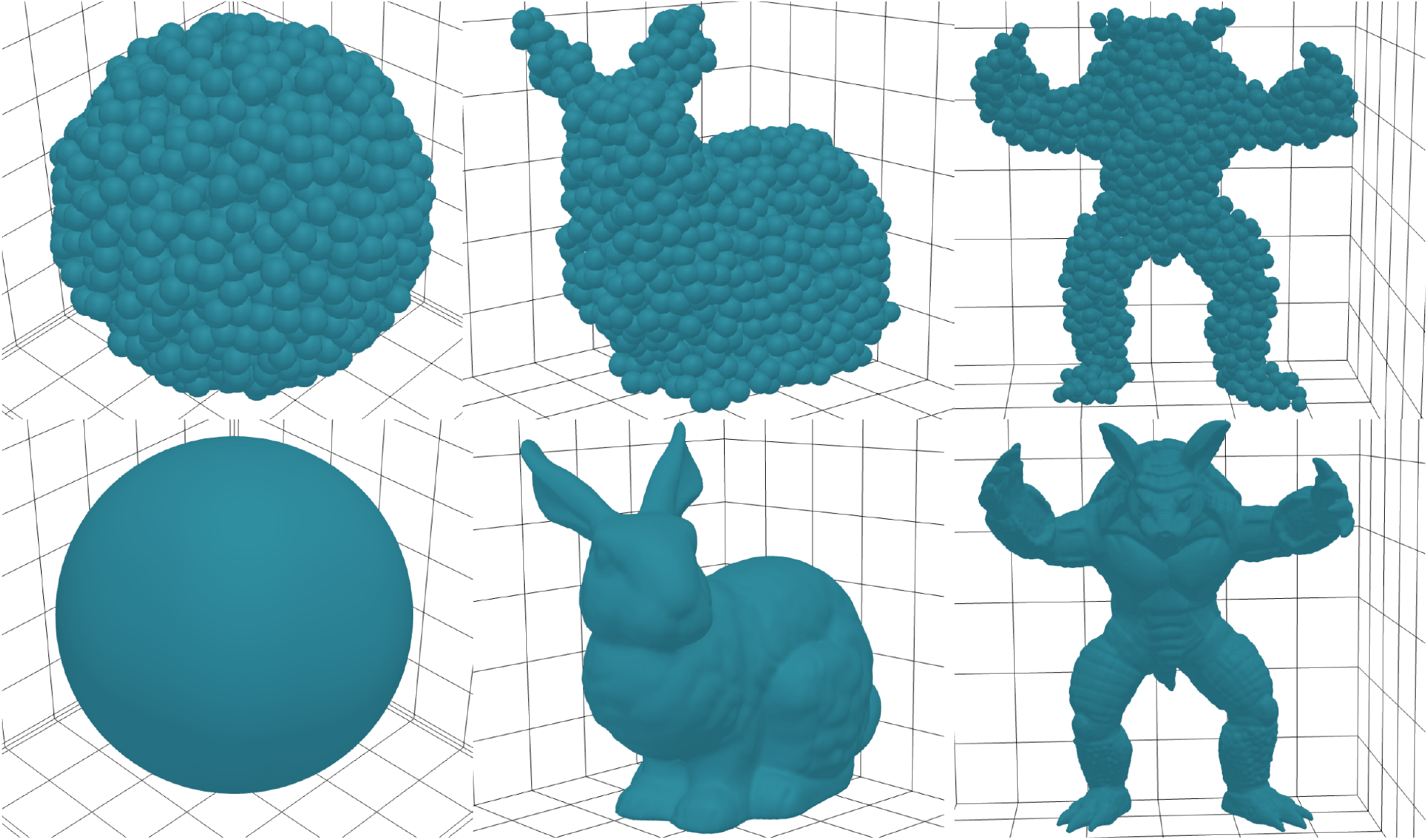
Mesh volume sampling examples with the data module. Top row images showcase spheroid samples with *N* = 2, 000 agents. Bottom row shows the original mesh.

**Supplementary Figure S5:**
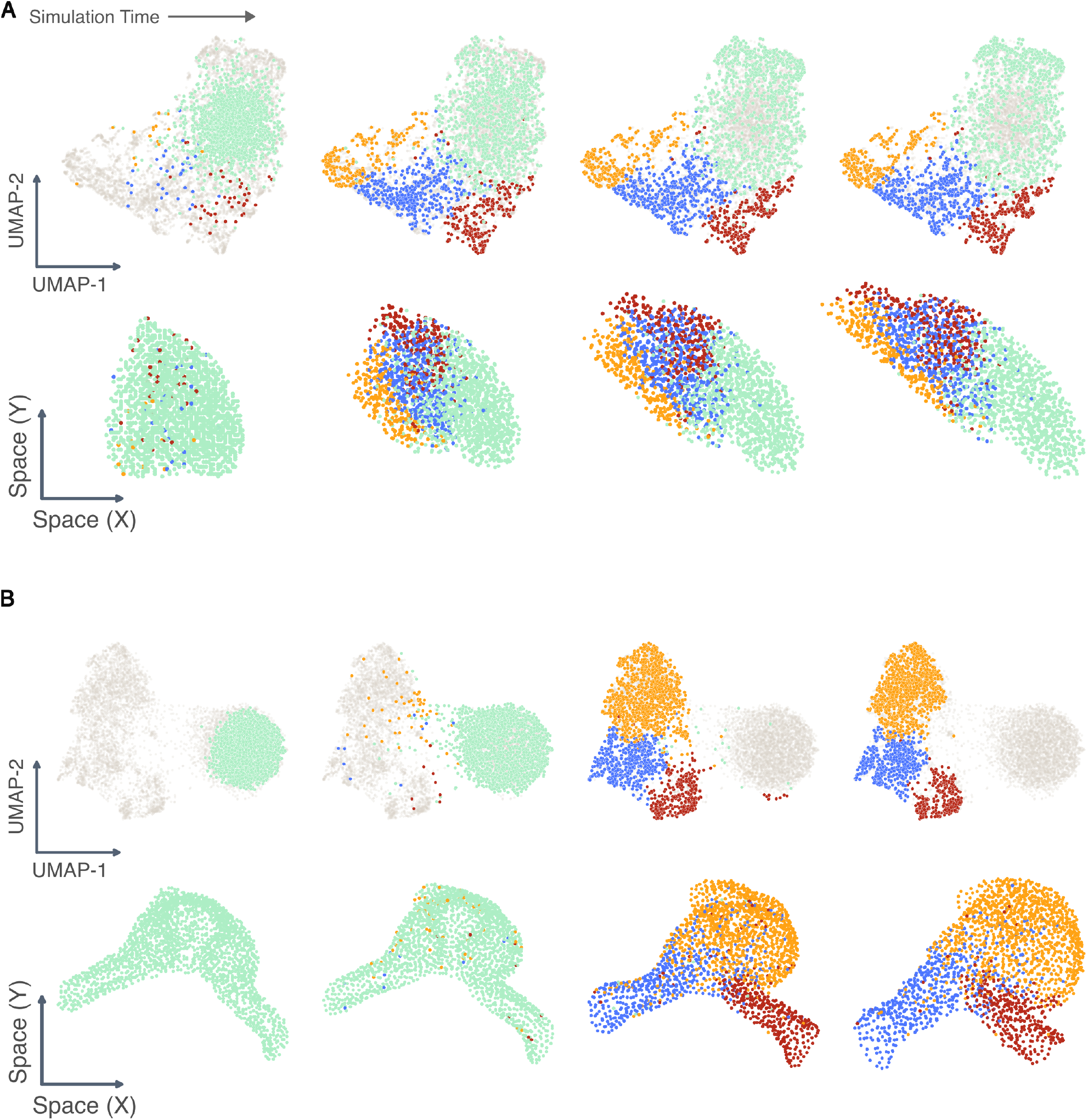
Quantification of molecular expression and spatial patterning for real data. **A, B**. Molecular expressions along the trajectory, clustered with Leiden and projected onto a 2D UMAP (top) and onto the XY plane (bottom) for the mouse limb [44] (A) and developing myocardium [45] datasets.

## Notes

https://github.com/Computational-Morphogenomics-Group/waxmorph

### Competing Interest Statement

The authors have declared no competing interest.

### Summary of Updates

completed missing acknowledgement sections; author affiliations updated;

